# Dentate Neurogenesis Modulates Dorsal Hippocampal Excitation/Inhibition Balance Crucial for Cognitive Flexibility

**DOI:** 10.1101/2023.02.22.529526

**Authors:** Haowei Li, Risako Tamura, Daiki Hayashi, Hirotaka Asai, Junya Koga, Shota Ando, Sayumi Yokota, Jun Kaneko, Keisuke Sakurai, Akira Sumiyoshi, Tadashi Yamamoto, Keigo Hikishima, Kazumasa Z. Tanaka, Thomas J. McHugh, Tatsuhiro Hisatsune

## Abstract

Adult neurogenesis endows the hippocampus with unparalleled neural plasticity, essential for intricate cognitive functions. However, the manner in which sparse newborn neurons (NBNs) modulate neural activities and subsequently shape behavior remains enigmatic. Utilizing a newly engineered NBN-Tetanus Toxin mouse model, we non-invasively silenced NBNs and confirmed their crucial role in cognitive flexibility, as demonstrated through reversal learning in the Morris water maze and the Go/Nogo task in operant learning. Pairing task-based functional MRI (tb-fMRI) with operant learning revealed a dorsal hippocampal hyperactivation during Nogo task, implying that hippocampal hyperexcitability might underlie the observed behavioral deficits. Notably, resting-state fMRI (rs-fMRI) revealed enhanced functional connectivity between the dorsal and ventral dentate gyrus following NBN silencing. Further exploration of PV+ interneurons and mossy cells’ activities highlighted NBN’s integral part in preserving the excitation/inhibition balance within the hippocampus. Our findings emphasize how the neural plasticity driven by NBNs extensively modulates the hippocampus, sculpting cognitive flexibility.

## Introduction

Organized memories and controlled forgetting help us navigate complex environments, preventing cognitive overload (*1, 2*). This balance largely hinges on executive functions, predominantly comprising working memory, inhibitory control, and cognitive flexibility (*3–5*). The hippocampus, an essential hub for higher-order cognitive functions like learning and memory, is deeply intertwined with these processes (*6, 7*). Current consensus postulates that neural plasticity is likely the principal neural mechanism supporting the hippocampus’s role as a center for learning and memory (*8, 9*). This plasticity spans multiple dimensions, including dendritic, axonal, synaptic, and circuit plasticity. Concurrently, adult neurogenesis, a captivating ability inherent to the hippocampus, not only endows it with heightened neural plasticity but also offers fresh insights into its function in learning, memory, and other advanced cognitive tasks.

Research has shown that persistent neurogenesis occurs in the hippocampal dentate gyrus of both animals and humans, at least extending into middle-aged individuals (*10, 11*). During their pivotal first 3-6 weeks, these newborn neurons (NBNs) exhibit unique physiological properties, marked by heightened excitability and plasticity (*12*). This equips them with the capability to recruit inhibitory GABAergic interneurons, positioning them as suppressors of the local mature granule cell activity in the dentate gyrus (*13, 14*). In hippocampal-dependent cognitive processes, various studies have substantiated the instrumental role of these NBNs, employing techniques such as tissue lesioning, X-ray ablation, and optogenetics, in the encoding of novelty, extinction of fear memories (*15, 16*), and the handling of spatiotemporal information (*17, 18*). As NBNs continuously integrate into existing neural circuits, advanced cognitive functions, including reversal learning(*19*), active forgetting(*20*), and pattern separation(*21*), are consequently realized (*22, 23*). Concurrently, the neural network restructuring triggered by the incorporation of NBNs can potentially destabilize prior memory circuits, catalyzing certain forms of forgetting (*24, 25*). However, this very mechanism provides a structural foundation for cognitive flexibility. Nevertheless, due to the sparse distribution of NBNs and limited observation tools (*26*), targeted modulation remains elusive, curtailing not only the unveiling of their distinctive roles but also a comprehensive description of their interactions within the hippocampus and the broader brain network (*27*). As we deepen our understanding of the functional heterogeneity exhibited along the hippocampal longitudinal axis (*28, 29*) and the physiological impacts of emotional stress on NBNs (*30, 31*), prior interpretations of the presence and functional significance of NBNs may need revision. Moreover, data regarding the contribution of NBNs to the excitation/inhibition (E/I) balance in the dentate circuitry is notably scant (*32*). Consequently, a pressing challenge lies in visualizing the contributions of NBNs to the hippocampal network in the most physiologically authentic conditions, offering an intuitive portrayal of their functionality, operational periods, and spheres of influence.

In this study, by combining features from two transgenic systems (*33–36*), we developed a novel transgenic mouse model, NBN-TeTX (newborn neuron-tetanus toxin expressing), that can non-invasively silence the functionality of NBNs. Through multiple reversal learning tasks (*26, 37*), we confirmed the impairments in inhibitory control and cognitive flexibility induced by silencing NBNs. Concurrently, task-based fMRI visually demonstrated the impact of silencing NBNs on the activation patterns of the hippocampus during tasks (*38, 39*). By integrating resting-state fMRI functional connectivity analysis with investigations into the activity of parvalbumin-positive interneurons and mossy cells in the dentate gyrus under various conditions (*40, 41*), we postulate that NBNs play an integral role in modulating the E/I balance among the dorsal dentate gyrus and contribute to the functional segregation between the dorsal and ventral hippocampus. These findings offer insights and a foundation for further elucidating the contributions of the hippocampus and NBNs to spatiotemporal perception and cognitive flexibility.

## Results

### NBN-TeTX Mouse Model for Non-Invasive Silencing of Newborn Neurons

Our primary focus is on the unique properties of adult-born neurons (3-6 weeks old) in memory, learning, and cognition. Thus, we have developed a tri-transgenic mouse model named NBN-TeTX (NBN, Fig. 1A, Table S1). In this model, administering tamoxifen to adult mice specifically induces the activation of CreERT2 only in adult-born stem/progenitor cells that express Nestin, without affecting neural stem cells during the mouse’s early developmental stage. As the labeled adult-born stem cells mature and start expressing αCamKII, TeTX is activated. It then inhibits the fusion of neurotransmitter vesicles with the presynaptic membrane by cleaving the synaptic vesicle protein VAMP2, thereby suppressing neurotransmitter release associated with the activity of newborn neurons (NBNs) during learning, memory, and cognitive tasks.

**Fig. 1.**
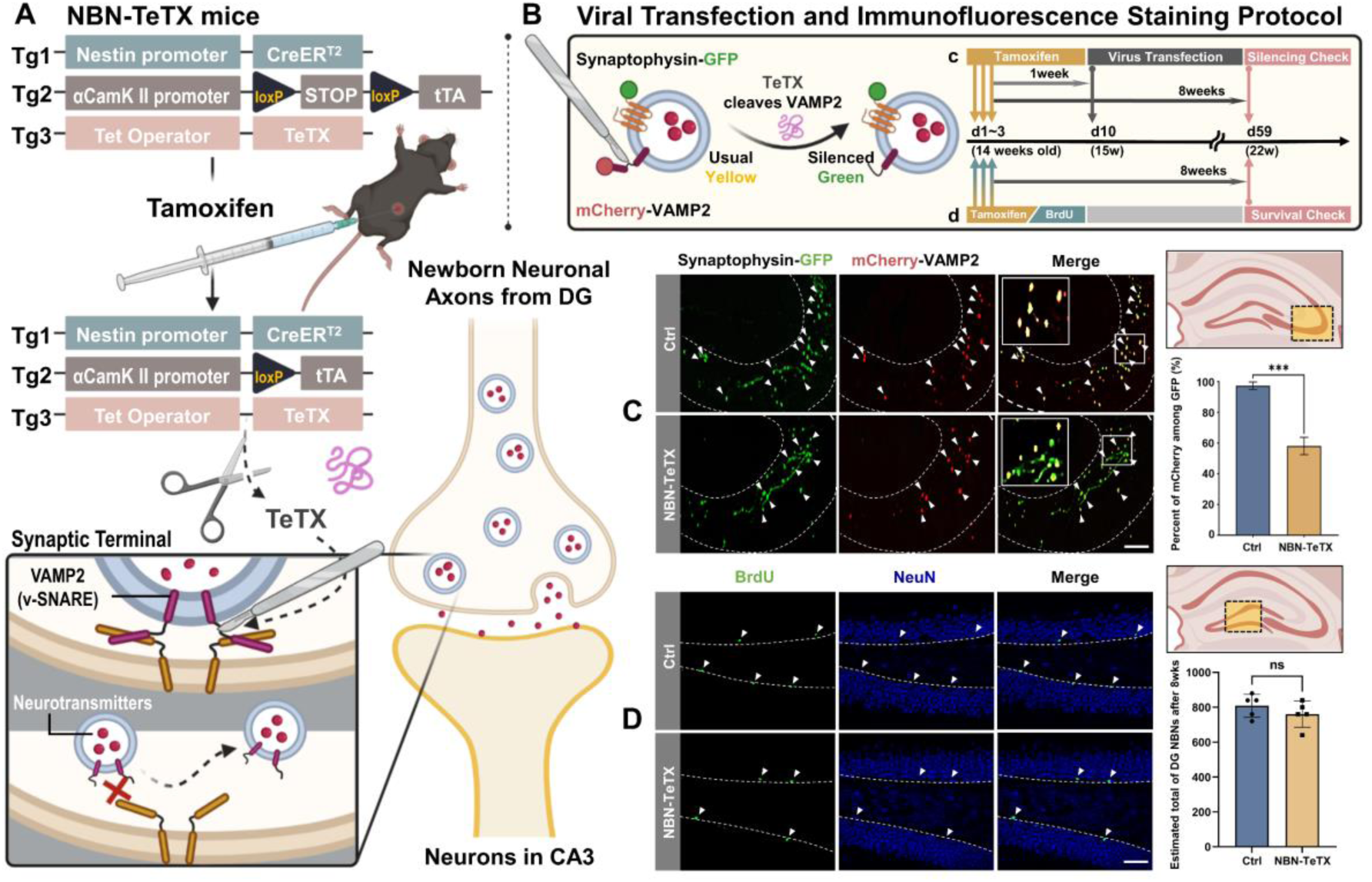
NBN-TeTX Mouse Model for Non-Invasive Silencing of Newborn Neurons. **(A),** NBN-TeTX mice are transgenic, carrying three recombinant genes. TeTX inhibits newborn neuron functionality by cleaving synaptic vesicle VAMP2 at neural terminals, thus restricting neurotransmitter release. **(B),** Fluorescent vector viral transfection method to validate the silencing rate. Synaptic vesicles at terminals are labeled with co-expressed Synaptophysin-GFP and mCherry-VAMP2. A week post-tamoxifen cessation, retroviral injections are given, with perfusion and staining after 8 weeks. Typically, co-localized VAMP2 and Synaptophysin in virus-transfected terminals display an overall yellow appearance due to combined mCherry (red) and GFP (green). In terminals with TeTX, VAMP2 cleavage separates mCherry from the synaptic vesicle membrane, resulting in only green GFP fluorescence. **(C),** In the CA3 area of NBN mice, ∼40% of NBN synaptic terminals showcase dislodged mCherry, in contrast to nearly none in the control group. Arrows highlight synapses co-fluorescing Synaptophysin-GFP and mCherry-VAMP2. (Ctrl: mean=97.265±1.382, NBN: mean=58.003±3.366, n = 3 per group; ***p<0.001, unpaired t-test; slice thickness: 40 µm, scale bar: 100 µm). **(D),** Post 8 weeks of tamoxifen administration, BrdU/NBN co-immunofluorescence staining indicates the mature NBN count in the NBN-TeTX group isn’t significantly different from the Ctrl group. Arrows pinpoint hippocampal dentate gyrus NBNs co-stained with BrdU/NeuN. (Ctrl: Average=733±32, NBN: mean=690±34, n = 5 per group, unpaired t-test; slice thickness: 40 µm, scale bar: 50 µm).

Using virus-mediated immunofluorescence staining, we confirmed the expression of TeTX in synaptic terminals of NBNs (Fig. 1B). In the NBN mouse model, we achieved a functional silencing of approximately 40% of NBNs. Specifically, after administering tamoxifen to 14- week-old adult mice and waiting one week for TeTX to be fully expressed and accumulate in the synaptic terminals of labeled NBNs, we then proceeded with retroviral injections to respectively label Synaptophysin (GFP) and VAMP2 (mCherry). Seven weeks post-viral injection (i.e., when NBNs are around 8 weeks old and fully matured), we conducted tissue slicing and immunofluorescence staining. Results indicated that, in the CA3 – the only projection area of the dentate gyrus NBNs – about 40% of the synaptic terminals in NBN mice showed residual green fluorescence of Synaptophysin-GFP due to the shedding of mCherry-VAMP2, compared to the control group. In contrast, synaptic terminals in the control group were almost entirely of a normal yellow color (Fig. 1C).

Furthermore, to ascertain if the NBN mouse model solely induced functional inhibition of NBNs without affecting their survival, we counted the NBNs in another set of experiments targeting the subgranular zone of the dentate gyrus. Slicing and co-staining with BrdU/NeuN immunofluorescence were done 8 weeks post-simultaneous tamoxifen and BrdU injection. Results depicted no significant difference in the number of surviving mature NBNs between the NBN group and the control group (Fig. 1D).

Consequently, we confirmed that the NBN-TeTX mouse model offers a non-invasive functional silencing window from the neonatal to mature phase for NBNs. By adjusting the administration timing of tamoxifen, this model facilitates temporally specific functional silencing of NBNs without compromising early developmental stage neural stem cells.

### NBN Silencing Impairs Spatial Reversal Learning

We employed a reversal learning task in the Morris water maze experiment to ascertain the impact of silencing NBNs on cognitive flexibility (Fig. 2A). Thirteen NBN-TeTX mice (NBN) and eleven control mice (Ctrl) received tamoxifen injections at 14 weeks of age, and three weeks later underwent 5 days of acquisition learning (D1-D5) followed by 5 days of reversal learning (D6-D10). Spatial probe tests were scheduled accordingly on Day 5 (after acquisition learning), Day 10 (after reversal learning), and Day 13 (Fig.2B). Throughout the 10-day training period, there was no significant difference between the two groups in swimming speed (Fig. 2D). As shown in Fig. 2C (D1-D5), there was no significant difference between the NBN and Ctrl groups in escape latency during the first 5 days of acquisition learning. In the first spatial probe test (P1), both groups showed no significant difference in the time spent in the target quadrant (NE), indicating that silencing mature NBNs did not exhibit significant difference in initial spatial learning (Fig. 2E, P1).

**Fig. 2.**
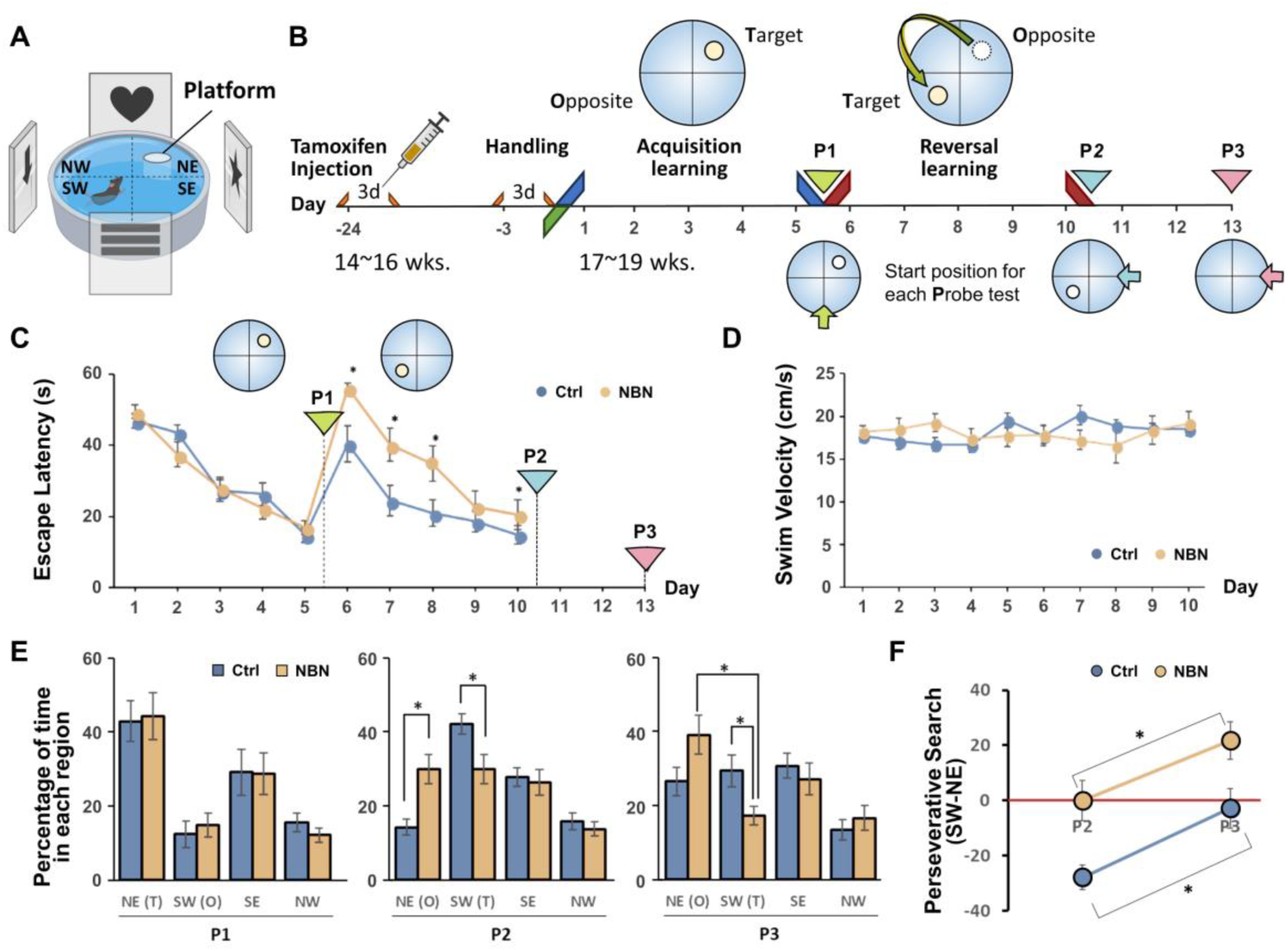
Impaired Spatial Reversal Learning During Morris Water Maze. **(A)**, Morris Water Maze (MWM) experimental setup. **(B)**, MWM experiment protocol. Probe tests P1 and P2 occurred immediately after the learning periods on days D5 and D10, respectively. P3 was scheduled for day 13. **(C)**, Escape latency progression (in seconds) over the 10-day training. Significant differences between the groups were observed only during the reversal learning phase (D6-10: *p < 0.05; 2-way ANOVA (Bonferroni); mean ± SEM). **(D)**, No significant difference in movement speed (cm/s) was found throughout the 10-day training (two-way ANOVA). **(E)**, Percentage of time both mouse groups spent exploring each quadrant during the spatial probe trial: At P1, exploration time in each quadrant showed no significant difference between the groups (NE: p=0.8808; two-way ANOVA). At P2, differences were seen in time spent in the NE and SW quadrants (NE: p = 0.0045, SW: p = 0.0285; *p < 0.05; 2-way ANOVA (Bonferroni)). At P3, only the time in the SW quadrant differed between groups (SW: p = 0.0232; 2-way ANOVA (Bonferroni)). Within the NBN group, significant differences arose between time spent in the NE and SW quadrants (Ctrl: p = 0.6394, NBN: p = 0.0015; *p < 0.05; 2-way ANOVA (Bonferroni)). **(F)**, Changing preference of both groups for the NE and SW quadrants, where the platform was located, from P2 to P3 (Ctrl: p = 0.0099, NBN: p = 0.0465; *p < 0.05; 2-way ANOVA (Bonferroni)).

During reversal learning (D6-D10), the platform was moved to the SW quadrant. Throughout the 5-day reversal learning process, the NBN group consistently displayed a significantly longer escape latency compared to the Ctrl group, although this gap gradually diminished (Fig. 2C, D6-D10). In the second spatial probe test (P2), the two groups of mice displayed differential area preferences. Ctrl mice explored the new location of the platform (SW) more, while the NBN group still favored the original platform location (NE) (Fig. 2E, P2). The third spatial probe test (P3) was conducted on Day 13 to assess the final long-term memories formed by both groups after 10 days of initial and reversal learning. At the P3 time point, Ctrl mice spent an equal amount of time exploring the previous two target locations (NE and SW), while the NBN group still displayed a pronounced preference for the NE quadrant (Fig. 2E, P3). By comparing the percentage search time for the NE and SW quadrants in P2 and P3 between the two groups, we observed a shift in preference in both groups. The Ctrl group transitioned from a preference for the platform’s initial NE quadrant to an equal preference for both quadrants where the platform had been located, while the NBN group maintained its attachment to the platform’s initial quadrant (Fig. 2F). These data suggest that the persistence in the initial memory after silencing mature NBNs led to the failure of spatial reversal learning in NBN mice, thereby impeding the realization of cognitive flexibility.

### NBN Silencing Impacts Inhibitory Control in Reversal Learning

To further confirm the effect of silencing NBNs on mice’s cognitive flexibility, we conducted an operant-learning experiment using the Go/Nogo paradigm. This was done to evaluate the mice’s inhibitory control by varying the conditions of reinforcement learning (Fig. 3A). Initially, we performed a head-plate fixation surgery on 11 NBN mice and 9 Ctrl mice aged 13-15 weeks and conducted a 3-day handling, initiating water restriction on the last day of the handling. This was followed by 3 days of light-lick training. The light cue was always on, and it briefly turned off when the mouse licked, immediately followed by a water reward. This established a connection between the light stimulus (Cue) and the water reward (Reward) before silencing the function of NBNs. No significant difference in the total number of licks was observed in the 5-minute post-training test (Fig. S2A). Tamoxifen injections were then administered to mice aged 14-16 weeks and, three weeks later, a 10- day Go task and a 10-day Nogo task began (Fig. 3B).

**Fig. 3.**
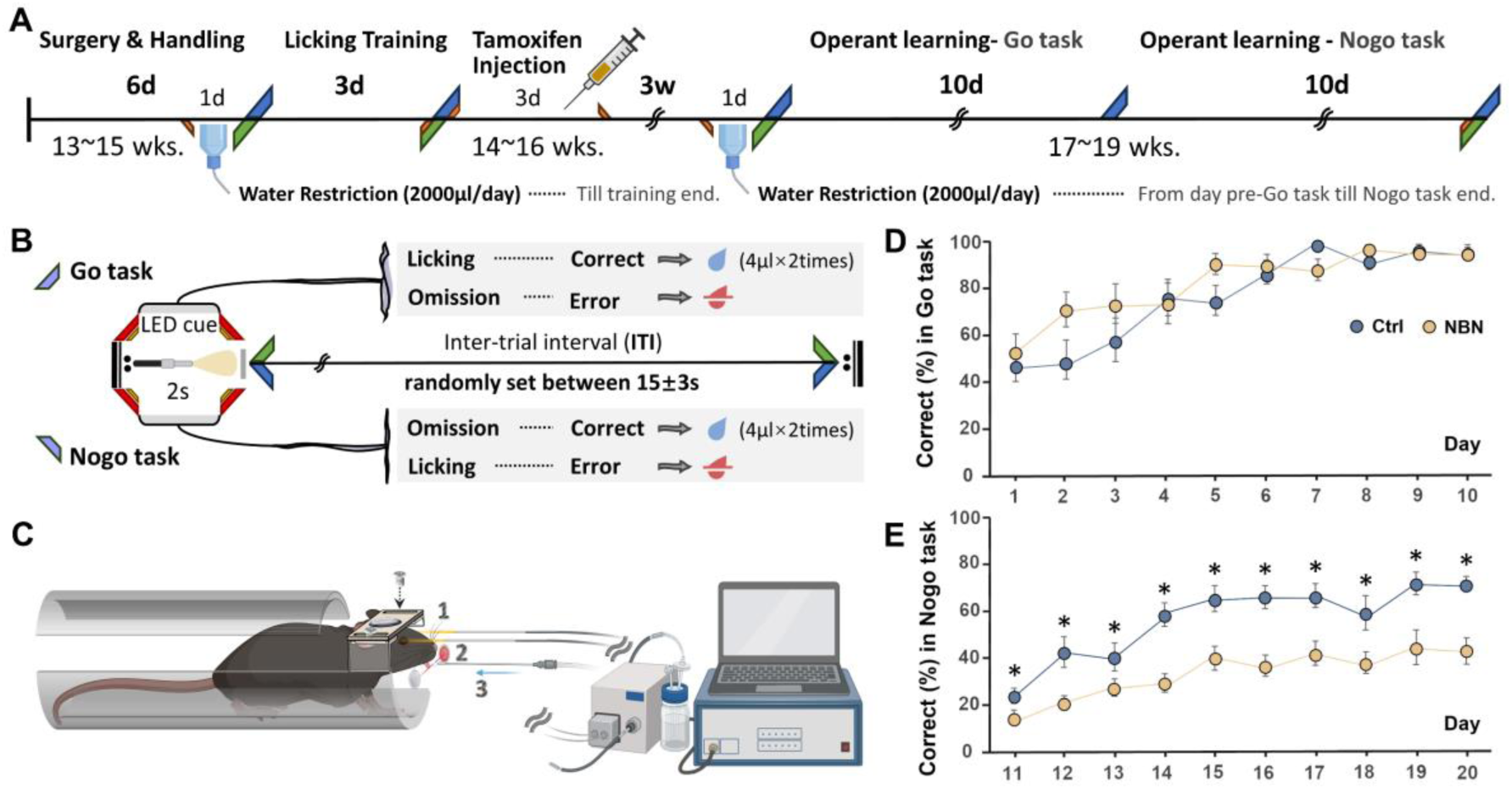
Impacted Inhibitory Control in the Nogo Task of Operant-Learning. **(A)**, Operant learning protocol based on light cues: 11 NBN and 9 Ctrl mice participated. A 3-day light reflex training preceded tamoxifen administration to ensure consistent light cue-water reward conditioning across both groups. **(B)(C)**, Illustrate rule settings and schematics for Go and Nogo tasks. Mice are secured on a custom operant learning bed via a pre-attached head plate. **(D)**, Represents each group’s accuracy trend across the 10-day Go task. By day 10, both groups consistently achieved over 95% accuracy, showing no marked inter-group differences. (Day 9 accuracy: Ctrl: 96.37%, NBN: 94.96%; Day 10: Ctrl: 95.72%, NBN: 95.68%; two-way ANOVA; Ctrl: n = 9, NBN: n = 11; mean ±SEM). **(E)**, Shows accuracy trends for each group during the 10-day Nogo task. The NBN group’s accuracy consistently trailed the Ctrl group throughout. (Genotype: F (1, 18) = 16.6545, p = 0.0007; *p < 0.05; 2-way ANOVA (Bonferroni); Ctrl: n = 9, NBN: n = 11; mean ±SEM).

During the D1-D10 Go task (Fig. 3B, Go task), both the NBN and Ctrl groups of mice gradually improved their accuracy rate (Correct%) of correct licking over the 10 days, reaching around 95% by the end of the task (Fig. 3D). During this period, there was no significant difference in the increasing trend of accuracy between the two groups. The daily total number of licks (Fig. S2B), the number of incorrect licks before reaching 100 correct licks daily (Fig. S2C), and the daily response latency (reaction speed to the light stimulus) (Fig. S2D) also showed no significant differences between the groups.

Subsequently, during the D11-D20 Nogo task (Fig. 3B, Nogo task), the mice were required to inhibit their learned impulse to lick water during the light stimulus and wait for the light cue to end to receive the water reward. We observed that throughout the 10-day Nogo task, the NBN group displayed a behavioral inhibition defect in suppressing licking. Despite not receiving a water reward, they frequently licked water during the light cue, causing their accuracy rate to consistently lag that of the Ctrl group (Fig. 3E). This suggests that, in reinforcement learning tasks with changing conditions, silencing NBNs appears not to affect the learning of initial rules. However, it significantly impacts inhibitory control, making new reinforcement learning challenging or requiring more extended periods to achieve.

### Hippocampal Activation Shifts in Initial and Reversal Learning via fMRI

To further understand the functional impact of the silencing of NBNs on the hippocampal activation state, we first combined the operant-learning experiment with functional magnetic resonance imaging (fMRI). By using task-based fMRI (tb-fMRI), we analyzed the activity state of the hippocampus of awake mice in real-time and non-invasively during the Go/Nogo tasks.

Six NBN and six Ctrl mice (aged 13-15 weeks) underwent head plate fixation surgery. Apart from being trained in the fMRI device with a 90db SE-EPI scanning sound to acclimate them to the environment, all other procedures were as described above (Fig. 4A). On the first day of the Go task and the first day of the Nogo task, we scanned the two groups of mice with fMRI (6 min), with specific task rules as shown in Fig. 4B. Based on hippocampal anatomical structures, we set up a hippocampal mask and conducted an ROI analysis. We assessed hippocampal activity changes post light cue (2-4 seconds) against baseline (average of the last 4 seconds per trial).

**Fig. 4.**
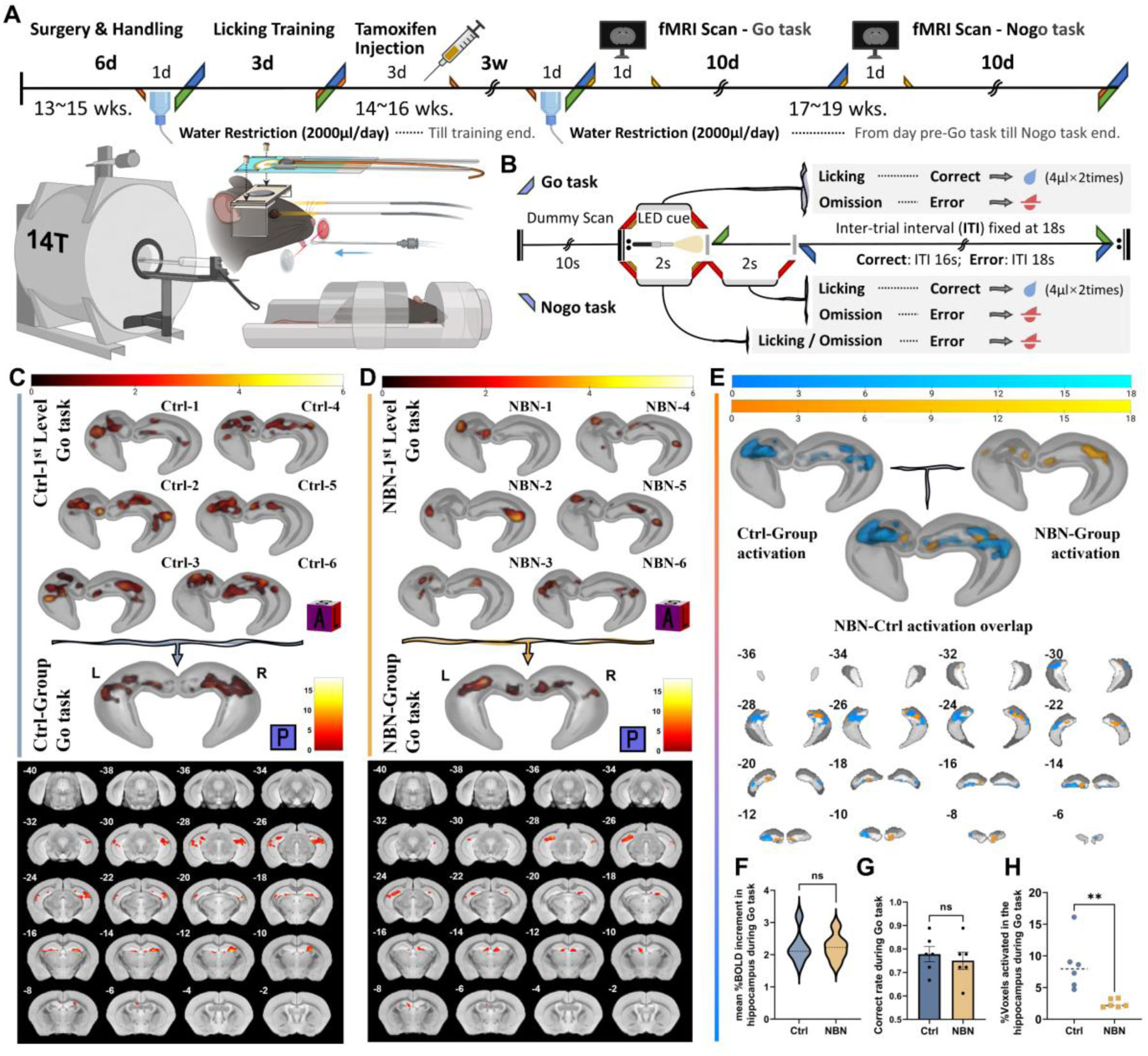
Hippocampal Activation in Go task via fMRI. **(A)**, Operant-learning protocol with light cues inside fMRI captures 18 trials over 6 minutes on day one of Go and Nogo tasks. **(B)**, Go and Nogo task rules for fMRI: 20-second trials. In Nogo, post-light cue licking grants a reward, countering freezing due to fMRI conditions. 2-second after the light cue aligning with the setting of active condition in the fMRI analysis, which used a hippocampal anatomical mask. **(C)**, 3D rendering of hippocampal activity in the Ctrl group’s individual and intra-group Go task (pFDR<0.05); posterior coronal visualization details section’s Bregma relation. 2nd-level: Max T value: 15.25, cluster KE=448. **(D)**, 3D rendering of hippocampal activity in the NBN group’s individual and intra-group Go task (pFDR<0.05); posterior coronal visualization details section’s Bregma relation. 2nd-level: Max T value: 14.29, cluster KE=216. **(E)**, Overlaid 2nd-level intra-group results for Ctrl and NBN during Go task highlight spatial activation differences. **(F)**, Both groups exhibit elevated %BOLD in hippocampus 2-4s post-light cue. No significant difference observed (Ctrl: 2.258, NBN: 2.236; n = 6 per group; two-tailed, unpaired t-test). **(G)**, Correct licking rate in fMRI-captured Go task trials: Ctrl: 0.7778, NBN: 0.7500; no significant group difference (n = 6 per group; two-tailed, unpaired t-test, mean ± SEM). **(H)**, Ratio of activated to total hippocampal voxels per individual calculated. Ctrl’s Go task activated voxel percentage significantly higher than NBN’s (Ctrl: 8.524%, NBN: 2.456%; **p<0.005, two-tailed, unpaired t-test; n = 6 per group).

In the Go task, the licking accuracy rate of both groups during the 18 trials in 6 minutes has no significant difference (Fig. 4G), however, the fMRI analysis results were intriguing. From the 1st-level analysis (individual) results of the hippocampal activation pattern of each individual in both groups, we observed similar activation patterns within the groups (Fig. 4C, D, top). By calculating the percentage of significantly activated hippocampal voxels, we confirmed that the activation range of the Ctrl group’s hippocampus was significantly larger than that of the NBN group (Fig. 4H). Through the 3D rendering of the 2nd-level intra-group analysis results (Fig. 4C, D, bottom, see also Fig. S3B) and the overlaid display of the activated voxel coronal montage (Fig. 4E), one can intuitively observe the spatial distribution differences in hippocampal activation. Surprisingly, the 2nd-level inter-group analysis did not return voxels with significant differences. We further extracted the BOLD signal values throughout the scan and calculated their percentage change (%BOLD change). The results suggested that there was no significant difference in the BOLD intensity elevation level during the Go task between the two groups. These results indicate that silencing NBNs during the Go task did not cause significant differences in hippocampal activity levels and behavioral performance but did result in a sharp reduction in spatial distribution.

However, on the first day when the rule switched to the Nogo task, the activation patterns of the two groups underwent a dramatic reversal. First, similar to the results of the Nogo task in the operant-learning experiment conducted outside of the fMRI, during the initial 6 minutes covered by the fMRI scan, which consisted of 18 trials, although neither group could achieve a high accuracy rate, the licking accuracy rate of the Ctrl group was still higher than that of the NBN group (Fig. 5F). Subsequently, in the individual analysis, we observed dissimilarity in the spatial distribution of hippocampal activation between the two groups (Fig. 5A, B, left). The activation range of the Ctrl group’s individuals during the Nogo task was significantly smaller than that of the NBN group and was smaller than their own range during the Go task. The NBN group, on the other hand, displayed a larger hippocampal activation range than its own performance during the Go task. Further, the intra-group analysis highlighted the dissimilarity between the two groups ’ hippocampal activities (Fig. 5A, B, right, see also Fig. S3B). Subsequently, in the inter-group analysis, we confirmed a significant difference in hippocampal activity between the NBN and Ctrl groups (Fig. 5C). Notably, whether looking at the spatial distribution of intragroup results (Fig. 5G) or the spatial distribution of intergroup differences, it seems to be mainly located in the dorsal region of the hippocampus.

**Fig. 5.**
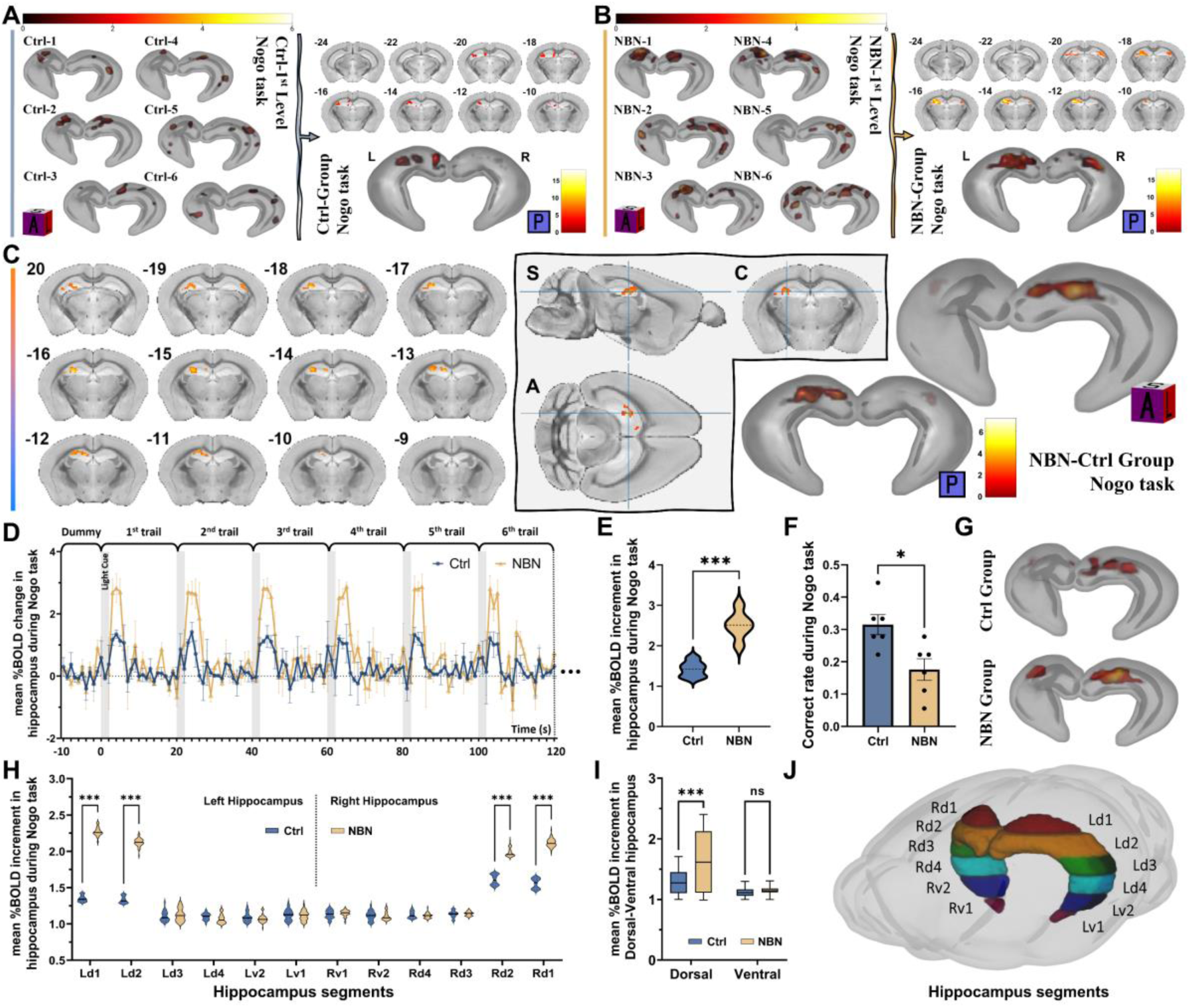
Hippocampal Activation in Nogo task via fMRI. **(A)**, 3D rendering of hippocampal activity in the Ctrl group’s individual and intra-group Nogo task (pFDR<0.05). **(B)**, 3D rendering of hippocampal activity in the NBN group’s individual and intra-group Nogo task (pFDR<0.05). **(C)**, 2nd-level inter-group analysis results for the Nogo task (NBN>Ctrl). The cluster with the peak T-value contains 372 voxels, peak T-value: 5.28 (pFDR<0.05). **(D)**, Hippocampal %BOLD signal changes are plotted over the initial 2-min frame during Nogo task (mean ± SD, n = 6 per group). **(E)**, NBN group exhibit significantly elevated %BOLD in hippocampus against the Ctrl group within 2-4s post-light cue (***p<0.001, two-tailed, unpaired t-test). **(F)**, In the Nogo task, the NBN group’s accuracy rate in the initial 18 trials was significantly lower than the Ctrl group (*p<0.05, two-tailed, unpaired t-test, mean ± SEM). **(G)**, 3D rendering of the 2nd-level intra-group analysis during the Nogo task. **(H)**, BOLD signal measurements from various layers of the hippocampus reveal notable intensity changes, especially in the NBN group’s d1 and d2 layers (***p<0.001, 2-way ANOVA (Bonferroni)). **(I)**, Evaluations of the average BOLD intensity changes between the dorsal and ventral hippocampus indicate predominant dorsal activation in the NBN group during the Nogo task (Dorsal: ***p<0.001, 2-way ANOVA (Bonferroni)). **(J)**, An illustration demarcates the mask used for BOLD signal extraction. The hippocampus is segmented into layers, with a breakdown of the dorsal (d1-d4) and ventral (v1-v2) regions. Both sides of the hippocampus are assessed independently.

To speculate on the reason for the activity differences concentrated in the dorsal hippocampus, we first calculated the percentage change in BOLD intensity of the two groups during the Nogo task. The results showed that within 2 seconds after the end of the light cue, during the peak period of the BOLD signal, the NBN group was significantly higher than the Ctrl group (Fig. 5D, E). Then, we divided the hippocampus along the longitudinal axis into 6 layers (Fig. 5J). The upper 2/3 (4 layers) were considered the dorsal hippocampus (d1-d4), and the lower 1/3 (2 layers) was considered the ventral hippocampus (v1-v2). We then extracted the BOLD signals from each layer of the bilateral hippocampus and calculated the percentage change. The results showed that during the Nogo task, the dorsal hippocampal layers d1-d2 of both groups contributed mainly to the rise in the BOLD signal, but the NBN group exhibited a significantly higher activation relative to Ctrl. However, no significant differences were observed in the entire ventral hippocampus between the two groups (Fig. 5H, I). This difference was observed bilaterally in the hippocampus (Fig. 5H).

In summary, tb-fMRI successfully visualized the transition in hippocampal activity patterns during the Go/Nogo task. On the first day of the Go task, prominent activity observed in both mouse groups indicated hippocampal engagement in detecting novelties and initiating learning. However, NBN silencing led to a significant reduction in the spatial distribution of activated regions. Even though no behavioral or mean %BOLD intensity change differences were observed in the NBN group during the Go task, the reduced activation areas in conjunction with poorer subsequent Nogo task performance indicated changes in initial learning strategies and intensities due to impaired NBN function. In the Nogo task, the Ctrl group completed the new rule learning with considerably less hippocampal activity compared to the heightened neural activity seen in the NBN group. Notably, the primary differences were observed in the dorsal hippocampus.

### NBN Silencing Enhances Functional Connectivity Between Dorsal and Ventral Dentate Gyrus

To understand the heightened dorsal activation in the NBN group during the Nogo task and the effect of NBN silencing on the hippocampal functional network, we conducted a 6- minute resting-state fMRI scan on 14 NBN mice and 14 Ctrl mice each, three weeks post-tamoxifen injection while they were under anesthesia.

In ROI-ROI functional connectivity analysis, we divided the hippocampus into the dentate gyrus (DG), CA1, and CA2&3, defining 6 VOIs based on dorsal (d) and ventral (v) sections (Fig. 6A). In the 2nd-level intra-group analysis, we observed differences in the hippocampal functional connectivity patterns between the NBN and Ctrl groups. Notably, in the Ctrl group, there was significant negative connectivity between dDG-vDG and dCA2&3- vCA2&3, which was absent in the NBN group. The functional connectivity difference between dDG-vDG persisted in the subsequent 2nd-level inter-group analysis (Fig. 6B). To quantify this disparity, we utilized Fisher’s z-transformation to convert Pearson correlation coefficients into z-values. As shown in Fig. 6C, for ROIs either within the dorsal or ventral areas, functional connections were predominantly positive with no significant difference between the two groups. However, for cross-region connections between dorsal and ventral, both groups displayed negative functional connectivity, but the NBN group exhibited a general trend of reduced negative values. Specifically, there was a significant difference between the NBN and Ctrl groups in dDG-vDG correlations (Fig. 6D).

**Fig. 6.**
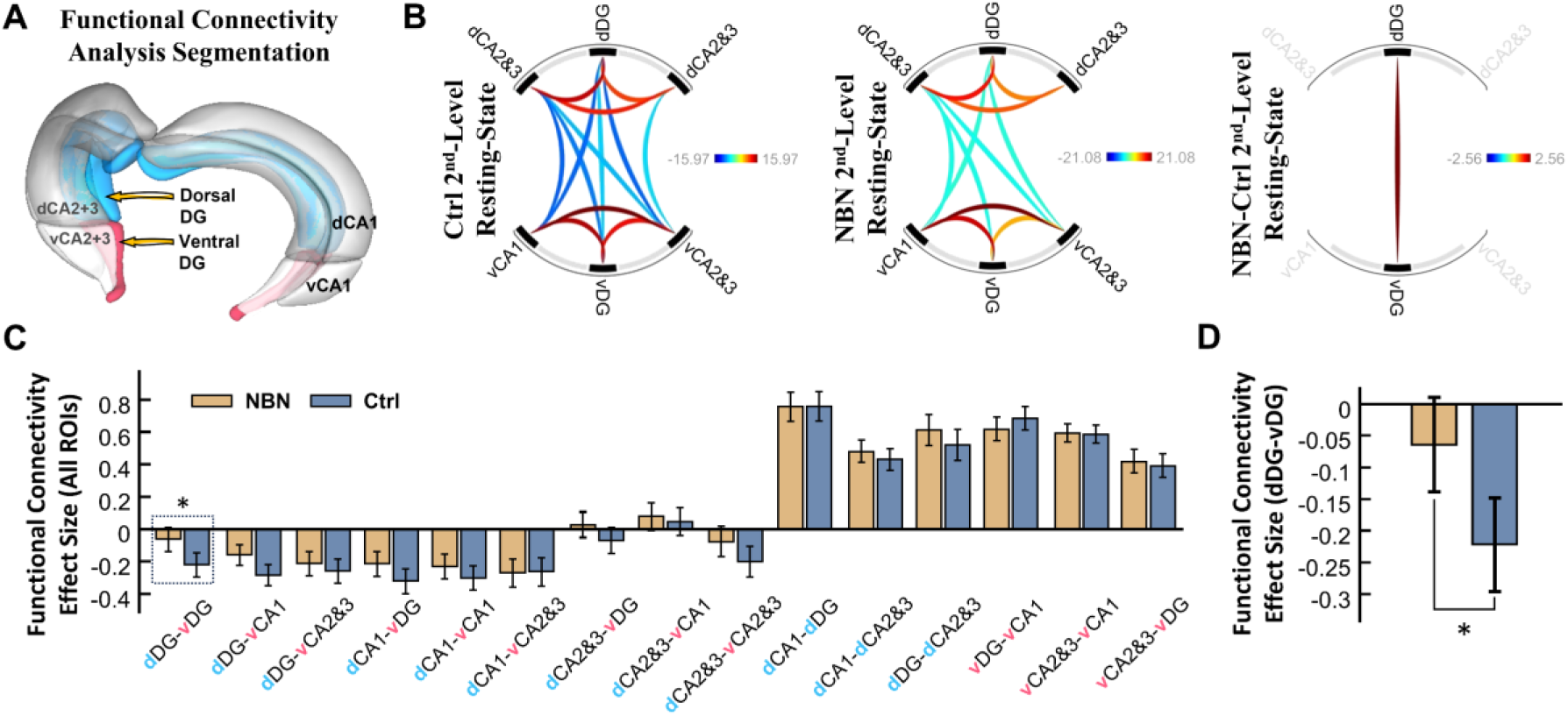
Enhanced Functional Connectivity Between Dorsal and Ventral Dentate Gyrus Post-NBN Silencing. **(A)**, Schematic of ROI settings for the resting-state fMRI functional connectivity analysis. The bilateral hippocampus is segmented based on anatomy and dorsal-ventral distinctions: dDG (upper 2/3), vDG (lower 1/3), dCA1, vCA1, dCA2&3, and vCA2&3. **(B)**, From left to right: 2nd-level intra-group functional connectivity for the Ctrl and NBN groups, 2nd-level inter-group analysis, and a chord diagram for NBN > Ctrl (T (26) =2.56, FDR-corrected p<0.05 ROI-level threshold (ROI mass/intensity), n = 14 per group). **(C)(D)**, Comparison of z-values (Fisher-transformed) between the groups. A notable difference in functional connectivity between NBN and Ctrl was seen in dDG-vDG (beta=0.16, T (26) =2.56, *p=0.016, two-sample t-test, mean ±SEM; n = 14 per group).

Our findings suggest that silencing the NBNs altered the functional connectivity patterns between the dorsal and ventral regions of the hippocampus. Typically, negative functional connectivity implies that as one region becomes more active, the other becomes less active. However, in the NBN group, this negative connectivity between the dorsal and ventral dentate gyrus was notably reduced, leading to an increase in synchronous activity between these regions. Such synchronicity indicates stronger or more enhanced functional connectivity, which, in this context, seems to be an abnormal consequence of NBN silencing.

### Dynamic Changes in the E/I Balance of the Dentate Gyrus Following NBN Silencing

Hyperactivation of the dorsal hippocampus in the NBN group during reversal learning hints at changes in the Excitation/Inhibition (E/I) balance. Hyperactivation of the dorsal hippocampus in the NBN group during reversal learning hints at changes in the Excitation/Inhibition (E/I) balance. To explore this, we examined c-Fos expression in various cell types, including NBNs, inhibitory component parvalbumin-positive GABAergic interneurons, and excitatory component mossy cells, in the hippocampus’s dentate gyrus after the Nogo task (Fig. 7A, B). We also assessed this under post-Go task and resting state conditions.

**Fig. 7.**
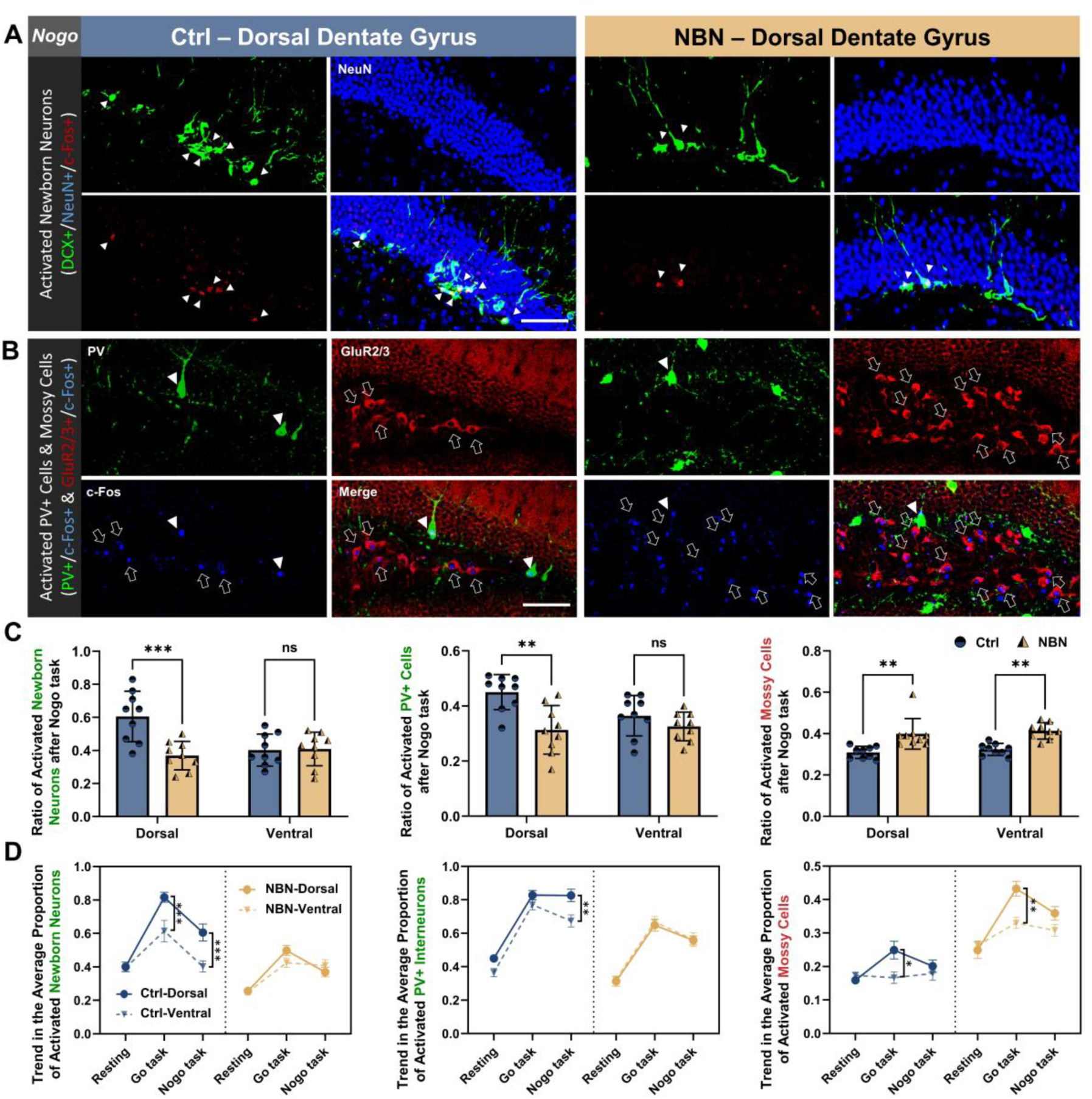
Newborn Neurons Modulates the Activity of DG E/I Components. **(A)**, Displays immunofluorescence of NBNs activity (DCX+/NeuN+/c-Fos+) post-Nogo task. The ratio of cells exhibiting c-Fos expression to the overall count of relevant cells in the dentate gyrus of each slice was determined (n = 3 per group, Scale bar: 100 µm). **(B)**, Shows the immunofluorescence of parvalbumin-positive GABAergic interneurons (PV+/c-Fos+) and mossy cells (GluR2/3+/c-Fos+) activity post-Nogo task (n = 3 per group, Scale bar: 100 µm). **(C)**, Indicates c-Fos+ NBNs, PV+ interneurons, and mossy cell proportions in the hippocampal dentate gyrus post-Nogo task for both groups (*p<0.033, **p<0.002, ***p<0.001, 2-way ANOVA (Bonferroni), mean ± SEM). **(D)**, Details dorsal-ventral changes and trends in activated NBNs, PV+ interneurons, and mossy cells in the hippocampal dentate gyrus under each condition (NBN: n = 3; *p<0.033, **p<0.002, ***p<0.001, 2-way ANOVA (Bonferroni), mean ±SEM; n = 3 per group at each condition).

Post-Nogo task, the NBN group displayed a significant drop in c-Fos+ NBNs (DCX+/NeuN+) in the dorsal dentate gyrus compared to the Ctrl group. However, this difference was not present ventrally (Fig. 7A, C; see also Fig. S4A, B). Additionally, in the NBN group, the number of c-Fos+ PV+ interneurons in the dorsal region was reduced compared to the Ctrl group. Meanwhile, the expression of c-Fos+ in mossy cells surpassed that of the Ctrl group in both dorsal and ventral areas (Fig. 7B, C; see also Fig. S4A, B).

After the Go task, the NBN group had fewer c-Fos+ NBNs in both regions compared to the Ctrl group. In the resting state, NBN activity in the NBN group remained consistently lower across both regions than in the Ctrl group. c-Fos+ PV+ interneuron activity in the NBN group showed a decline dorsally in the resting state and post-Go task. Conversely, c-Fos+ mossy cell increased across both domains.

As experiments progressed from resting to the Nogo task, NBN activity during tasks increased relative to the resting phase for both groups. Yet, this increase was always less pronounced in the NBN group (Fig. 7D). Post-Nogo task in the Ctrl group, dorsal PV+ interneuron activity was higher than in the ventral area, but this was not the case in the NBN group. Mossy cell activity in the NBN group remained consistently elevated across both dorsal and ventral hippocampus. This data suggests the altered PV+ interneuron activity in the absence of excitatory NBN inputs might explain the observed BOLD changes in the dorsal hippocampus during the Nogo task.

## Discussion

Undeniably, the unique contribution of newborn neurons (NBNs) to cognitive functions and behavioral regulation is far more impressive than their sparse numbers would suggest. Adult hippocampal neurogenesis, while widespread in mammals, including humans, remains controversial mainly due to the restricted techniques for modulation and observation in humans (*42, 43*). Previous research highlighted the role of NBNs in novelty encoding, spatial navigation, and learning, primarily by manipulating their numbers or using X-ray irradiation to disrupt them (*23, 44–47*). However, Inconsistencies in these findings (*48, 49*) underscored the need for a non-invasive method to modulate NBN numbers without altering the hippocampal anatomy. Recognizing these challenges, our study employed the newly-engineered NBN-TeTX tri-transgenic mouse model to provide clarity. By integrating the Nes-CreERT2 and TetO-TeTX systems, we achieved targeted and temporal modulation of NBNs (*33, 35*). Through our method, we managed to silence around 40% of NBN functions for up to 8 weeks post a single tamoxifen injection, without affecting overall neurogenesis levels or mature newborn neuron numbers.

The dentate gyrus (DG), as the hub of neurogenesis, plays a pivotal role in hippocampal-dependent tasks, notably pattern separation (*50–52*). Our research contributes to a growing body of evidence that underscores the role of NBNs in differentiating highly similar situations (*21, 53*). Pattern separation inherently requires the discrete encoding of similar data, necessitating sparse activity patterns within the DG (*54*). NBNs, compared to their mature dentate granule cell counterparts, may provide transient support for this sparse activity during specific phases (*55*). We propose that NBNs modulate hippocampal activation patterns, potentially influencing the DG’s excitation/inhibition (E/I) balance.

Diving deeper into the roles of hippocampal subregions, our research adds nuance to the ongoing debate surrounding the distinct functionalities of the dorsal and ventral parts of the hippocampus, particularly in relation to cognitive flexibility (*56–58*). Both our team and others are in the process of establishing experimental systems to gather fMRI data from mice engaged in the Go/Nogo tasks (*37, 38*). Our fMRI findings indicate that when NBN function is compromised, the hippocampus exerts heightened effort, manifesting in a broader or heightened activation pattern, to accommodate and revise new memories. Such shifts not only alter initial learning strategies and intensities, but also disrupt the typical connectivity within the hippocampus, which could potentially be the essence of compromised cognitive flexibility during reversal learning. Intriguingly, we noted an upregulation when observing the functional connectivity between the dorsal and ventral hippocampus. This increased involvement of the ventral hippocampus introduces an added complexity, potentially acting as an interference during reversal learning and amplifying the activation observed in the dorsal region. Hence, NBNs might maintain functional separation and network stability within the hippocampus by fine-tuning its internal E/I balance.

To unravel the potential underlying mechanisms, our analysis associates the functional impairment of NBNs with altered activities of PV+ inhibitory interneurons and mossy cells, fundamental components of the DG makeup (*59*). The parvalbumin-expressing (PV+) interneurons possess dense dendritic structures and form synapses extensively with excitatory cells across various layers. By receiving ample excitatory impulses and monitoring excitation levels, these PV+ cells distribute potent inhibitory signals uniformly to surrounding neurons, playing a crucial role in maintaining the E/I balance within neural circuits (*41, 60*). Meanwhile, mossy cells in the hilus, serving as the first to project glutamate inputs to newborn neurons (*61*), are considered sentinels of the DG network (*62*). Their broad projection also allows them to regulate the E/I balance within the DG’s granule cells (*63, 64*). Moreover, this regulation arises not only from providing direct excitatory glutamate signals to granule cells but also from projecting to a series of GABAergic interneurons, indirectly supplying inhibitory signals to granule cells. Our study reveals that the functional impairment of NBNs is associated with a concomitant reduction in the activity of PV+ inhibitory interneurons within the dorsal hippocampus during reversal learning. These insights intimate the NBNs’ pivotal role in harmonizing the balance of local neural circuits, influencing both excitatory and inhibitory neural populations (*40, 65*). The dorsal hippocampus’s observed hyperactivity during reversal learning further elucidates these complexities (*40, 61*). In the context of fully functional 3-6-week-old NBNs, their excitatory stimulation towards surrounding PV+ cells ensures that these GABAergic interneurons exert a robust inhibitory effect across the entire dentate gyrus (DG), guaranteeing sparse activation within the DG during reversal learning. However, when NBN functionality is compromised, the diminished recruitment of GABAergic interneurons leads the PV+ cells into a dormant state. Consequently, this results in hyperactivation of the DG during reversal learning, thereby hindering cognitive flexibility (Fig. 8).

**Fig. 8.**
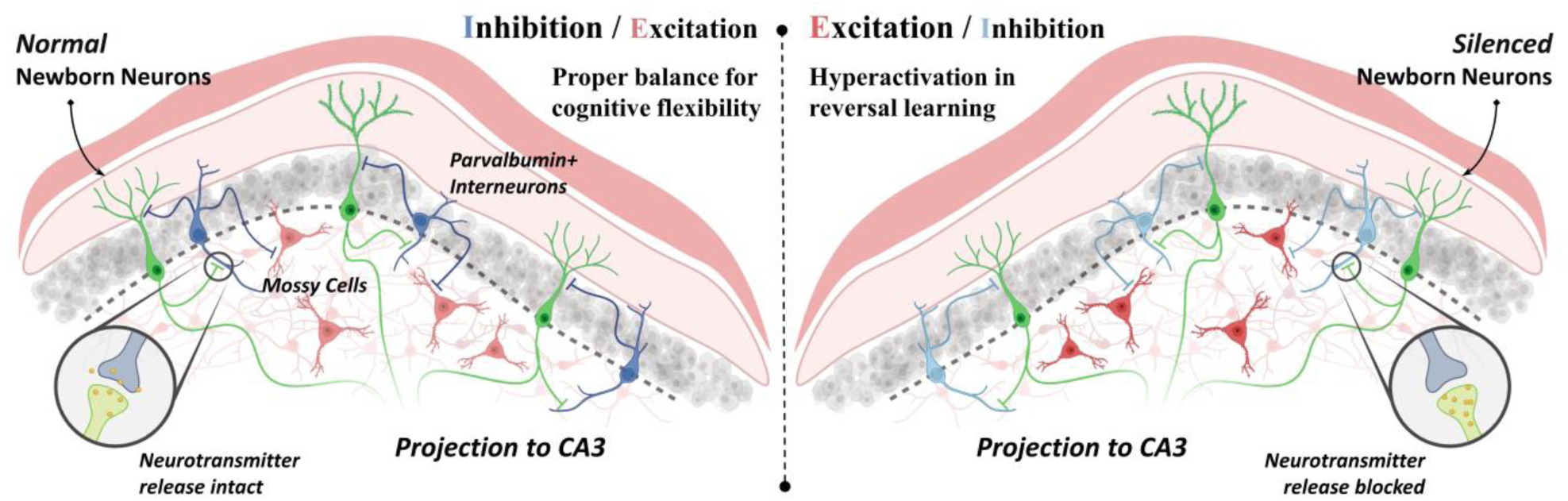
E/I Imbalance in the Dentate Gyrus Following NBN Silencing. With functional NBNs, the robust inhibitory input from dorsal dentate gyrus PV+ cells ensure moderate hippocampal activation during cognitive flexibility tasks. When NBN function is compromised, reduced PV+ cell activity results in hippocampal hyperactivation, accompanied by increased mossy cell activity.

A broader perspective reveals the symbiotic relationship between working memory and inhibitory control in fostering cognitive flexibility. The hippocampus, alongside the prefrontal cortex (PFC), is increasingly recognized as central to cognitive flexibility, evident in reversal learning tasks (*23, 66–68*). Such insights prompt considerations of neurodegenerative diseases like Alzheimer’s disease, temporal lobe epilepsy, or post-traumatic stress disorder, where hippocampal damage is a hallmark. Typically, by the time these conditions are diagnosed, significant hippocampal structural alterations, such as volume reductions and sclerosis, are evident. Hence, there’s a pressing need for methodologies that can specifically emulate the cognitive and behavioral aberrations resultant from hippocampal damage, facilitating investigations into potential etiologies and laying the groundwork for preemptive identification and risk assessment prior to overt structural damages. Our results intimate that employing fMRI to gauge the intensity of hippocampal activation and changes in longitudinal functional connectivity could be a promising avenue for the early detection and evaluation of the state of hippocampal neurogenesis and cognitive deficits.

In summation, our research, through the NBN-TeTX mouse model, elucidates the NBNs’ significance within the dentate gyrus circuitry. In concert with GABAergic interneurons and mossy cells, they collaboratively preserving the hippocampus’s dorso-ventral functional segregation and the excitation/inhibition balance, laying the foundation for cognitive flexibility.

Nonetheless, certain limitations warrant attention. Translating findings from a mouse model to humans always requires caution. Moreover, as fMRI provides indirect neural activity insights, the interplay between NBNs and hippocampal cell types warrants further investigation. Future research should explore NBNs’ overarching effects across neural systems and their evolution with age and external influences.

## Materials and Methods

### Experimental Design

The primary objective of our study is to elucidate the role of newborn neurons (NBNs) in modulating the overall functionality of the hippocampus, with a particular focus on their contribution to cognitive flexibility. We will employ the NBN-TeTX mouse model, wherein NBNs are to be non-invasively silenced. This model allows us to probe potential changes in neural activities and consequent behaviors.

1. NBN Silencing using the NBN-TeTX Model: By utilizing the NBN-TeTX mouse model, we aim to non-invasively silence the NBNs, which will pave the way to uncover the inherent intricacies of NBN functionalities.
2. Behavioral Assessment: We will monitor the behavioral control by NBNs during initial and reversal learning phases using the Morris water maze and operant-learning tasks.
3. fMRI Analysis: In adherence to our non-invasive approach, fMRI will be employed to characterize shifts in hippocampal activation patterns during operant-learning post-NBN silencing. This analysis involves observing and comparing during both initial learning and reversal learning phases.
4. Cellular Activity Analysis in the Dentate Gyrus: By employing immunohistochemistry, we aim to analyze the activity patterns of key cellular components within the dentate gyrus.

### Transgenic composition of NBNs-TeTX mice

The NBN-TeTX (newborn neuron-tetanus toxin expressing) triple-transgenic mouse model was constructed by combining two distinct transgenic systems: Nes-CreERT2 System (*35*) and TetO-TeTX System (*33*). For mice bearing these three recombinant genes, TeTX expression can be specifically induced post tamoxifen injection, suppresses neurotransmitter release by cleaving VAMP2 located in synaptic vesicles, thereby non-invasively inhibiting NBN function in the NBN-TeTX mice. For detailed mechanism, see Supplementary Materials.

### Genotyping

To ascertain the presence of the target recombinant gene in transgenic mice, genotypes were identified via PCR, utilizing DNA extracted from mouse tails. Mice aged over three weeks had a 5 mm segment of their tails excised and subsequently placed in a lysis buffer mixture (100 mM Tris, 5 mM EDTA, 200 mM NaCl, 0.5% Tween-20, pH 8.5) containing proteinase K (Takara Bio Inc., Otsu, Japan). This mixture was incubated at 55 °C for at least 12 hours. The extracted DNA was then reconstituted in 50 μL of TE buffer and served as the PCR template. Detailed primer sequences, reaction solution compositions, and reaction conditions pertinent to the genotyping process are presented in Table S1.

### Tamoxifen administration

Tamoxifen was employed to induce Cre recombinase activity in CreERT2-expressing cells. It was combined with a solution of corn oil and 100% ethanol at a 9:1 ratio, resulting in a tamoxifen concentration of 30 mg/ml. The test tube, enveloped in aluminum foil, was incubated at 53 °C for an hour with periodic agitation to ensure tamoxifen dissolution. Subsequent incubation at 37 °C for a minimum of 30 minutes was conducted with intermittent shaking. This mixture was maintained at 37 °C until administered to the mice. Over a span of three days, mice were intraperitoneally administered a daily dose of 180 mg/kg. To calibrate the dosage, mice were weighed daily during the administration period. Sterilization of the needle tip was achieved using 70% ethanol before and after each dosing. For hygiene purposes, a fresh tamoxifen mixture was prepared daily. To negate potential tamoxifen influence on experimental outcomes, all experimental mice (including both NBN-TeTX and control groups) were injected with tamoxifen between 14 to 16 weeks of age. Moreover, all behavioral studies were initiated three weeks post-tamoxifen administration (i.e., when mice were between 17 to 19 weeks of age).

### BrdU injection

Concurrent with tamoxifen administration, BrdU (FUJIFILM, 027-15561) was introduced to assess any alteration in the quantity of surviving newborn neurons in NBN-TeTX mice 8 weeks subsequent to tamoxifen dosing. BrdU was prepared as a 10 mg/ml solution in saline and was delivered intraperitoneally at 100 mg/kg across three sequential days. For the discernment and enumeration of matured newborn neurons, immunohistochemical staining employing BrdU/NeuN markers was conducted.

### Retrovirus preparation and infection

The Moloney viral vector was generously donated by Professor Nakashiba (*34*). All vector solutions were prepared at a concentration of 1g/l in TE buffer (pH 8.0). Retroviral injections commenced three days post-tamoxifen administration. Under anesthesia, mice received an injection of the virus into the right dentate gyrus (DG) via a mineral oil-filled glass micropipette attached to a Nanoject II (Drummond) at coordinates (-2.06 mm AP, 1.25 mm ML, -1.75 mm DV) and (-2.7 mm AP, 2.0 mm ML, -1.75 mm DV). At each location, 897 nl (69 nl, 13 injections) of the retrovirus was administered at a rate of 0.8 L/min. Post-injection, the micropipette remained in situ for 3 minutes before extraction.

### Fixation, Sectioning, and Immunohistochemistry

Mice underwent transcardiac perfusion with PBS (composition in mM: 137 NaCl, 8.1 Na2HPO4, 2.68 KCl, and 1.47 KH2PO4) followed by 4% paraformaldehyde (PFA). Brains, post-perfusion, were fixed overnight at 4°C in 4% PFA, then underwent a gradient dehydration over three days in 10%, 20%, and 30% sucrose solutions. Subsequently, brains were embedded in O.C.T. Compound (Sakura, Tokyo, Japan) and sectioned into 40 μm slices, utilizing a cryostat (Thermo Fisher Scientific). Free-floating sections underwent immunofluorescence using standard protocols, post-washing with TBS (composition in mM: 137 NaCl, 2.68 KCl, 25 Tris, pH 7.2). A list of primary antibodies and their dilutions were as follows:

GFP/mcherry: Anti-GFP antibody [EPR14104] (1:2000, abcam, Cat# ab183734) and Living Colors® DsRed Polyclonal Antibody (1:200, Clontech). BrdU: A (1:200, abcam). DCX: doublecortin (C-18) goat polyclonal IgG (1:200, Takara Bio, Cat# 632496). NeuN: Ms X Neuronal Nuclei (1:1000, Millipore, Cat# MAB377). PV: Monoclonal Anti-Parvalbumin antibody produced in mouse (1:1000, Sigma, Cat# P3088), GluR2/3: Anti-Glutamate Receptor 2 & 3 antibody (1:500, Millipore, Cat# AB1506). c-Fos: rabbit and goat polyclonal IgG (1:200, Santa Cruz Biotechnology, Cat# sc-52 & Cat# sc-52-G).

### Immunofluorescence Image Acquisition and Processing

For GFP/mCherry co-staining quantification at CA3-projecting synapses, images were captured using an oil-immersion lens at 63x magnification on a confocal microscope (TCS SP2, Leica, Germany). Four slices per mouse were utilized for the synaptic co-staining count over the full dentate gyrus span (n = 3 per group). Additional imaging of immunolabeled specimens was done using confocal laser scanning microscopes (FLUOVIEW FV3000, Olympus, Japan) with 20x objectives.

To estimate the total NBN count (BrdU+/NeuN+) in the hippocampal DG, systematic sampling was employed. Three interval slices (each 40μm thick) covering the entirety of the DG were selected for each genotype (n = 5). Using a z-stack under a 40x objective, NBNs in both the left and right DG were counted for each selected slice. The cell count average from three sections per mouse was then multiplied by 60, based on the 2.4mm standard length of the DG in mice, to approximate the total cell count across the entire DG.

For c-Fos-associated immunofluorescence staining, resting state slices were procured from two mouse groups without any prior learning or training experiences. Following removal from the breeding room during light cycle, these mice were immediately perfused and fixed. Slices related to the Go and Nogo tasks came from mice perfused and fixed within 15 minutes post-completion of the learning tasks. In each case, three male mice from both groups were chosen. For each condition, slices were sectioned at 40μm thickness from hippocampal areas, illustrating typical dorsal and ventral DG structures. The c-Fos expressing cells’ percentage in every slice was determined relative to the total target cells. Imaging involved z-stacks at a pixel resolution of 1024 x 1024. Image processing was executed via Fluoview Software Module 3 and further refined with Fiji/ImageJ (*69, 70*), ensuring no selective adjustments or modifications to specific image segments.

### Selection of behavioral experimental subjects

In all the tests, mice that possessed the three previously mentioned genes made up the experimental group, while the control group consisted of mice that lacked one or both transgenes. Additionally, for the operant-learning trials done in the framework of fMRI, only male mice were used in both the NBN-TeTX and control groups. This decision was taken to eliminate potential discrepancies in BOLD signals and variations in behavioral outcomes that could arise from factors like brain size differences and variations in anxiety levels. Mice chosen for these behavioral tests were administered tamoxifen when they were between 14 and 16 weeks of age and were subsequently tested from weeks 17 to 19. All the behavioral trials were conducted during the light phase of the day. During the experiments, the experimenter was unaware of the mouse genotypes. The care and handling of all animals were in accordance with the guidelines set by the Animal Care and Use Committee of the Graduate School of Frontier Science at the University of Tokyo.

### Morris Water Maze

In an effort to understand the effects of neuronal functional inhibition on reversal learning capabilities, a training regimen spanning 10 days alongside three spatial exploration trials were carried out. These involved 13 NBN-TeTX mice and 11 control mice. The said spatial exploration trials took place on days 5, 10, and 13.

The time to reach the platform (latency) was calculated and used as a criterion of spatial memory to evaluate the mice’s learning. The first five days (days 1-5) were dedicated to acquisition learning. The platform was set in the NE quadrant, with learning sessions repeated six times daily. The first spatial exploration trial (P1) was initiated immediately after the fifth day of acquisition learning. Reversal learning was the focus of days 6-10. During this phase, the platform was repositioned to the SW quadrant, and mice were trained three times daily, with sequential randomization and 30 seconds between each training session. Spatial exploration trial P2 followed right after the tenth day’s reversal learning.

In total, three spatial exploration trials (P1, P2, and P3) were conducted. While P1 and P2 aimed to gauge memory retention across the respective learning stages, P3 (carried out after a 72-hour gap post-day 10) assessed the establishment of long-term memory post the entire 10-day learning period. All data from this experiment were collected and analyzed using the animal experiment analytical software, Smart 3.0 (by Panlab Harvard Apparatus). The detailed protocol can be found in the Supplementary Materials.

### Operant-learning experiment

The operant-learning experiment in this study used an operant-learning apparatus that uses light stimulation to cue mice to lick water and has been used to measure the executive function of the mice accurately. Mice are light-cued while their heads are immobilized in this device, and water is given as a reward when a sensor near their mouths detects a tongue-extending lick in response to the light stimulus (Fig. 3C). We used this device to perform Go Task and Nogo Task to investigate the effects of silencing NBNs on executive function.

On the last day of the handling, the mice were restricted to a maximum daily water intake of 2000μl until the end of the training. Following the handling, a three-day reflex-to-light training was conducted. The light cue was constantly on and was briefly turned off when the mice actively licked and were rewarded with water (4μl×2 times).

Three weeks after tamoxifen injections, a 10-day Go task was performed. Again, the mice were restricted to a maximum daily water intake of 2000μl on the last day of the handling until the end of the experiments. During Go task, the mice were given a light cue of 2s duration, and if the mice licked voluntarily during this 2s, they were considered correct and given a water reward. Immediately after the Go task, a 10-day Nogo task was performed, during which the mouse was also given a 2-s light cue, was considered correct if it did not lick during this period, and was given water rewards immediately after the light cue.

However, the rules of the operant-learning experiments performed in fMRI are slightly changed (Fig. 3A). First, each trial was set to 20 seconds due to the block design and given a light cue during the initial 2s (0-2s). Meanwhile, in the Nogo task, the rule was set to be considered correct only when the mouse actively licked within 2s after the light cue went off, to reduce the possibility of false correctness due to mouse inaction (omission) caused by the relatively cold environment and noise during the fMRI scan. That is, the water reward was given only when the subject actually comprehended the rule change and performed the correct licking behavior. The detailed protocol can be found in the Supplementary Materials.

### Task-based fMRI analysis

Equipment and Location: The task-based fMRI analysis was conducted at the Graduate School of Frontier Sciences, The University of Tokyo (GSFS). A high-precision 14-T small horizontal bore animal scanner from Jastec, Japan was used, paired with a mouse head adapted cryocoil and the ParaVision 5.0.1 software suite.

Sample: The subject pool consisted of 6 male NBN-TeTX mice and 6 control male mice, all of which had successfully completed the licking training.

Procedure: Mice were positioned with their custom resinous headplate (attached during previous surgeries) at the center of the cryocoil. The headplate installation and preprocessing steps prior to scanning were adopted from the work of Dr. Thi Ngoc Anh Dinh (*39*). The water tube’s position was adjusted to facilitate easy reward access for the mice. To ascertain the correct positioning of the mice, tripilot images were taken. tb-fMRI scanning lasted for 6min10s, with a preliminary 10-second window to stabilize the fMRI signal.

The specific parameters for the fMRI scans were: SE-EPI: TR=2000ms, TE=18ms, flip angle=90°, matrix size=64×64, FOV=17.2×17.2 mm^2^, 20 slices, interleaving sequence, 0.7mm slice thickness, no gap, BW=300 kHz. T2-weighted anatomical images: TR=800ms, TE=24ms, flip angle=90°, matrix size=256×256, FOV=17.2×17.2 mm^2^, 20 slices, interleaved sequence, 0.7mm slice thickness, no gap.

tb-fMRI Data Preprocessing: mainly processed using SPM12 (https://www.fil.ion.ucl.ac.uk/spm/), and the downsample Australian Mouse Brain Mapping Consortium (AMBMC) template (*71*) was used for coregister, mask creation, and result mapping (10 times and reorient of the coordinates occupied the midpoint of the anterior commissure; Dimensions: 68×131×50 voxels, voxel size: 0.15×0.15×0.15 mm).

The 1^st^-level analysis was conducted using a general linear model (GLM) analysis. Group-level results were derived using a one-sample t-test with a significance level set at cluster level pFDR < 0.05. For discerning statistically significant activations, a threshold was set at pFDR < 0.05 with a cluster size threshold 15 voxels. The detailed protocol can be found in the Supplementary Materials.

### Resting-state fMRI analysis

Location and Equipment: The rs-fMRI study was carried out at the Okinawa Institute of Science and Technology Graduate University (OIST). MRI data collection employed an 11.7-T small horizontal bore animal scanner (BioSpec 117/11, Bruker, Ettlingen, Germany), along with a mouse head-adapted cryocoil and ParaVision 6.0.1 software.

28 mice participated in this study, with 14 being NBN-TeTX mice and the remaining 14 as controls. Mice were anesthetized using isoflurane (3% for initiation and 1.5% during MRI). Each mouse was placed in a prone position on a custom MRI bed fitted with a bite bar and gas mask. Core body temperature was maintained at 37.0 ± 1.0 °C via a water-circulating pad, with monitoring facilitated by an MRI-compatible rectal temperature probe (Model 1025, SA Instruments). Tripilot images were taken to ensure correct mouse positioning in relation to the coil and magnet’s isocenter. The resting-state fMRI ran for 6 minutes, where the imaging parameters included:

SE-EPI: TR = 1800 ms, TE = 18 ms, flip angle = 90°, matrix size = 96 × 96, FOV = 20 × 20 mm^2^, 22 slices, slice thickness of 0.5 mm, no slice gap, BW = 500 kHz, and 200 total volumes. T2-weighted anatomical images: TR = 3374 ms, TE = 30 ms, flip angle = 90°, matrix size = 256 ×256, FOV = 20 ×20mm^2^, 34 slices, slice thickness of 0.5 mm, no gap, with 2 averages.

rs-fMRI Data Preprocessing: mainly processed using SPM12 and CONN toolbox (*72*) and 3D Slicer was used for the segmentation and definition of the regions of interest (ROI) (*73*). The detailed protocol can be found in the Supplementary Materials.

### Statistical Analysis

Statistical analyses for immunofluorescence staining and behavioral experiments were conducted using either unpaired t-test or two-way repeated measures ANOVA with Bonferroni post hoc test, as appropriate. GraphPad Prism 9.4.0 (673) for Windows was the primary tool for these analyses. For fMRI correlation analyses, the software’s built-in algorithm was employed. Significance was set at *P<0.05. Unless otherwise specified, error bars represent mean±SEM.

## Acknowledgments

We are grateful to Prof. Ryoichiro Kageyama and Prof. Itaru Imayoshi for their generous support of the mouse models needed for this experiment. We thank Dr. Toshiaki Nakashiba for the gift of Moloney virus. We also thank students Qiong Ding and Shuto Mine for their help in data collection. Some of the diagrams in the article were created with BioRender.com.

## Author contributions

Conceptualization: HL, HA, KZT, TJM, TH

Methodology: HL, RT, DH, HA, JKoga, SA, KS, AS, KZT, TJM, TH

Investigation: HL, RT, DH, JK, SA, SY, JKaneko, AS, TY, KH, TH

Visualization: HL, JKoga, KS, AS, TH

Supervision: HL, TJM, TH

Writing—original draft: HL, TJM, TH

Writing—review & editing: HL, TJM, TH

### Data and materials availability

All data available provided in the main text or the supplementary materials. Any additional information required to reanalyze the data reported in this paper is available from the lead contact upon request.

