## Supplementary Materials for "Dentate Neurogenesis Modulates Dorsal Hippocampal Excitation/Inhibition Balance Crucial for Cognitive Flexibility"

Haowei Li *et al.*

**This PDF file includes:**

Supplementary Text
Figs. S1 to S4
Tables S1

**Supplementary Text**

NBN Silencing Mechanism in NBN-TeTX Triple Transgenic Mice

The NBN-TeTX (newborn neuron-tetanus toxin expressing) triple-transgenic mouse model was constructed by combining two distinct transgenic systems:

1. Nes-CreERT2 System: This system facilitates the permanent labeling of neural stem/progenitor cells at specified time points by modulating the administration of tamoxifen. The gene Tg1 in this system expresses the Cre/ERT2 protein under the Nestin promoter (Nestin-Cre/ERT2). Nestin, an intermediate filament, is specifically expressed in neural stem and progenitor cells (*35*).
2. TetO-TeTX System: Here, tetanus toxin is expressed under the control of the αCamKII promoter. This toxin acts by cleaving VAMP2, subsequently inhibiting activity-dependent neurotransmitter release, thus silencing neuronal function. Within this system, Tg2 contains two loxP sequences downstream of the αCamKII promoter with a stop codon sandwiched between them. Once activated, it expresses tTA (tetracycline-controlled transactivator) downstream. Tg3, on the other hand, expresses tetanus toxin under the TetO (Tetracycline Operator) recombinant gene (TetO-TeTX) TetO-TeTX System: Here, tetanus toxin is expressed under the control of the αCamKII promoter. This toxin acts by cleaving VAMP2, subsequently inhibiting activity-dependent neurotransmitter release, thus silencing neuronal function. Within this system, Tg2 contains two loxP sequences downstream of the αCamKII promoter with a stop codon sandwiched between them. Once activated, it expresses tTA (tetracycline-controlled transactivator) downstream. Tg3, on the other hand, expresses tetanus toxin under the TetO (Tetracycline Operator) recombinant gene (TetO-TeTX) (*33*).

For mice bearing these three recombinant genes, TeTX expression can be specifically induced post tamoxifen injection, leading to inhibition of NBN function. Upon tamoxifen administration, its metabolite, 4-OHT (an estrogen analogue), binds to ERT. This binding allows Cre/ERT2 to translocate to the nucleus and exhibit Cre recombinase activity on Nestin-positive neural stem/progenitor cells. This action on the loxP of Tg2 results in the removal of the intervening STOP codon. When these modified stem/progenitor cells differentiate into neurons, the downstream αCamKII promoter becomes active, leading to the expression of tTA. This tTA subsequently binds to TetO in Tg3, initiating the transcription of TeTX. In its final action, TeTX suppresses neurotransmitter release by cleaving VAMP2 located in synaptic vesicles, thereby non-invasively inhibiting NBN function in the NBN-TeTX mice.

<u>Morris Water Maze Protocol Details</u>
In an effort to understand the effects of neuronal functional inhibition on reversal learning capabilities, a training regimen spanning 10 days alongside three spatial exploration trials were carried out. These involved 13 NBN-TeTX mice and 11 control mice. The said spatial exploration trials took place on days 5, 10, and 13.

The maze setup consisted of a 120 cm diameter pool filled with water, divided equally into four quadrants: NE, SW, SE, and NW. Hidden just below the water's surface (by 1 cm) was a 10 cm diameter platform that served as a refuge for the mice. The water's temperature was maintained between 23°C and 24°C. To obscure the underwater platform from the mice's sight, the water was tinted with an opaque, non-toxic white dye. Various cues in the form of distinct markings were affixed on the walls around the pool to help the mice orient themselves.

Over the course of the 10-day training span, which encompassed both acquisition and reversal learning stages, the mice were subjected to the following procedure during each session: Firstly, mice were randomly placed in one of the pool's quadrants, marking their starting point (Fig. S1A). The time to reach the platform (latency) was calculated and used as a criterion of spatial memory to evaluate the mice's learning. Upon locating the platform, the mice were allowed to remain there for 15 seconds, allowing them to memorize its position. If a mouse failed to locate the platform within a 60-second window, it was gently guided there with the aid of a wooden stick, and then allowed the 15-second memorization period. After the session, each mouse was dried with a towel before being returned to its cage.

The first five days (days 1-5) were dedicated to acquisition learning. Before these sessions, mice were handled for 5 minutes daily over three consecutive days to familiarize them with the experimenter. The platform was set in the NE quadrant, with learning sessions repeated six times daily. The sequences of these sessions were randomized to ensure an even spread across all quadrants. The first spatial exploration trial (P1) was initiated immediately after the fifth day of acquisition learning.

Reversal learning was the focus of days 6-10. During this phase, the platform was repositioned to the SW quadrant, and mice were trained three times daily, with sequential randomization and 30 seconds between each training session. Spatial exploration trial P2 followed right after the tenth day's reversal learning.

In total, three spatial exploration trials (P1, P2, and P3) were conducted. While P1 and P2 aimed to gauge memory retention across the respective learning stages, P3 (carried out after a 72-hour gap post-day 10) assessed the establishment of long-term memory post the entire 10-day learning period. In each probe test, the platform was removed from the pool, and mice were placed at any point in the pool, ensuring that all mice had the same starting point. All data from this experiment were collected and analyzed using the animal experiment analytical software, Smart 3.0 (by Panlab Harvard Apparatus).

Operant Learning Procedure Details

The operant-learning experiment in this study used an operant-learning apparatus that uses light stimulation to cue mice to lick water and has been used to measure the executive function of the mice accurately. Mice are light-cued while their heads are immobilized in this device, and water is given as a reward when a sensor near their mouths detects a tongue-extending lick in response to the light stimulus (Fig. 3C). This device delivers precise sensory stimulation by immobilizing the mouse's head, and this conditioned reflex is efficiently and successfully established because of the simple relationship between stimulus and response. We used this device to perform Go Task and Nogo Task to investigate the effects of silencing NBNs on executive function.

To fix the mouse's head during the experiments, a series of surgical procedures are required, including removing the hair on the top of the mouse's head after anesthesia, lifting the skin on the top of the skull and cutting it vertically, and making a small circular incision to expose the Bregma. The periosteum is infiltrated with a surface treatment agent for a few seconds before being pivoted to the sides to expose the skull. The Bregma is fitted with a customized resin fixation head plate, with the anterior edge of the head plate avoiding obscuring the mouse's eyes. The mice are returned to their home cage after the surgery and monitored until they are fully awake from anesthesia. The mice were returned to the breeding room and allowed to rest for one day before the three-day handling was taken, to familiarize them with the operators. On the last day of the handling, the mice were restricted to a maximum daily water intake of 2000µl until the end of the training. Following the handling, a three-day reflex-to-light training was conducted. The light cue was constantly on and was briefly turned off when the mice actively licked and were rewarded with water (4µl×2 times).

Three weeks after tamoxifen injections, a 10-day Go task was performed. Again, the mice were restricted to a maximum daily water intake of 2000µl on the last day of the handling until the end of the experiments. During Go task, the mice were given a light cue of 2s duration, and if the mice licked voluntarily during this 2s, they were considered correct and given a water reward. A 15±3-second inter-trial interval (ITI) was set between each trial. Immediately after the Go task, a 10-day Nogo task was performed, during which the mouse was also given a 2-s light cue, was considered correct if it did not lick during this period, and was given water rewards immediately after the light cue.

However, the rules of the operant-learning experiments performed in fMRI are slightly changed (Fig. 3A). First, each trial was set to 20 seconds due to the block design and given a light cue during the initial 2s (0-2s). Meanwhile, in the Nogo task, the rule was set to be considered correct only when the mouse actively licked within 2s after the light cue went off, to reduce the possibility of false correctness due to mouse inaction (omission) caused by the relatively cold environment and noise during the fMRI scan. That is, the water reward was given only when the subject actually comprehended the rule change and performed the correct licking behavior.

Task-based fMRI analysis protocol

tb-fMRI Data Preprocessing: mainly processed using SPM12 (https://www.fil.ion.ucl.ac.uk/spm/), and the following steps, illustrated in Fig. S3A, were undertaken:

1. Image reconstruction and Format Conversion: Images were exported in DICOM format and then transformed into NIfTI format using cm2niix in MRIcroGL. This step also entailed slice timing correction.
2. Image reorientation: Images were manually rotated and flipped so that the coordinate origin was positioned at the mid-point of the forebrain comma, aligning it with the standard reference space.
3. Voxel resizing: The nominal voxel size was magnified tenfold using SPM12 to streamline the image processing.
4. Initial image exclusion: The first 10s of EPI images from each scan was discarded.
5. Realignment: The EPI images underwent motion correction, realigning them to the mean using SPM12. Images with a translation of more than 0.1 mm and a rotation of more than 1 degree were excluded from the analysis.
6. Coregister: The T2 anatomical images were employed to co-register the EPI images with the Australian Mouse Brain Mapping Consortium (AMBMC) template (*71*). The template's dimensions and specifications were adjusted to ensure compatibility (10 times and reorient of the coordinates occupied the midpoint of the anterior commissure. Dimensions: 68×131×50 voxels, voxel size: 0.15×0.15×0.15 mm)
7. Normalization: EPI images were normalized using the downsample AMBMC template, and resliced to 0.15x0.15x0.15mm isotropic voxels. The number of hippocampal voxels was specifically noted at 13,965 for subsequent analysis of calculating the percentage of hippocampal activated voxels as presented in Fig. 4H.
8. Smoothing: EPI images were smoothed to a full width at half maximum of 0.3 mm.
9. 1st-level Analysis: A multiple regression analysis was employed to account for and remove the influence of six head motion parameters, along with the mean signals from white matter and ventricles. Use hippocampal anatomical appearance mask to focus the analysis inside the hippocampus.

<u>Resting-state fMRI analysis protocol</u>

Location and Equipment: The rs-fMRI study was carried out at the Okinawa Institute of Science and Technology Graduate University (OIST). MRI data collection employed an 11.7-T small horizontal bore animal scanner (BioSpec 117/11, Bruker, Ettlingen, Germany), along with a mouse head-adapted cryocoil and ParaVision 6.0.1 software.
Sample: 28 mice participated in this study, with 14 being NBN-TeTX mice and the remaining 14 as controls.
Procedure: Anesthetization and Positioning: Mice were anesthetized using isoflurane (3% for initiation and 1.5% during MRI). Each mouse was placed in a prone position on a custom MRI bed fitted with a bite bar and gas mask. Core body temperature was maintained at 37.0 ± 1.0 °C via a water-circulating pad, with monitoring facilitated by an MRI-compatible rectal temperature probe (Model 1025, SA Instruments). Tripilot images were taken to ensure correct mouse positioning in relation to the coil and magnet's isocenter. The resting-state fMRI ran for 6 minutes, where the imaging parameters included:
SE-EPI: TR = 1800 ms, TE = 18 ms, flip angle = 90°, matrix size = 96 × 96, FOV = 20 × 20 mm$^2$, 22 slices, slice thickness of 0.5 mm, no slice gap, BW = 500 kHz, and 200 total volumes. T2-weighted anatomical images: TR = 3374 ms, TE = 30 ms, flip angle = 90°, matrix size = 256 × 256, FOV = 20 × 20mm$^2$, 34 slices, slice thickness of 0.5 mm, no gap, with 2 averages.

rs-fMRI Data Preprocessing: mainly processed using SPM12 and CONN toolbox (*72*).

1. Image Reconstruction and Format Conversion: The image was first reformed, exported in DICOM, and then shifted to NIfTI format using dcm2niix in MRIcroGL. This step also entailed slice timing correction.
2. Image Reorientation: The image was manually rotated and translated to position the coordinate origin at the anterior commissure's midpoint, approximating the standard reference space.
3. Voxel Resizing: The nominal voxel size was inflated tenfold through SPM12 to optimize image processing.
4. Initial Volume Exclusion: The first 10 EPI volumes were discarded.
5. Motion Correction: EPI image motion was corrected using SPM12.
6. EPI Template Creation: A study-specific template was derived from the mean of the EPI images with SPM12.
7. T2 Template Creation: Another study-specific template was crafted using the T2 anatomical images via SPM12.
8. Coregister: The EPI template underwent linear and non-linear coregistration to the T2 template, and then to the previously detailed downsampled AMBMC.
9. Normalization: EPI images were standardized to the atlas template and resampled into 100 isotropic voxels.
10. Smoothing: EPI images were smoothed, targeting a full width at half-maximum of 0.2 mm.
11. Noise and Signal Removal: Six head movement parameters, combined with the average signals from white matter, cerebrospinal fluid, and ventricles, were eradicated using multiple regression analysis in CONN.
12. Temporal Filtering: EPI images underwent temporal filtration between 0.01 and 0.1 Hz with CONN.
13. Correlation Computation: CONN was utilized to compute and plot the correlation coefficient between each Region of Interest (ROI).
14. Template ROI definition: 3D Slicer was used for the segmentation and definition of the regions of interest (ROI) (*73*).

**A** Starting positions for mice in each trial of the Morris water maze training sessions (Acquisition learning, Reversal learning)

| Day | | Trial 1 | Trial 2 | Trial 3 | Trial 4 | Trial 5 | Trial 6 | Probe test |
|---|---|---|---|---|---|---|---|---|
| 1 | | NE | SE | NW | SW | SE | NW | |
| 2 | | NW | SE | NW | SE | SW | NE | |
| 3 | Acquisition | SE | NE | NW | SW | SE | NW | |
| 4 | (target: NE) | SE | NW | SE | SW | NE | NW | |
| 5 | | NE | NW | SE | NW | SE | SW | P 1 |
| 6 | | NE | NW | SE | | | | |
| 7 | | SE | NW | NE | | | | |
| 8 | Reversal | NE | SE | NW | | | | |
| 9 | (target: SW) | NW | NE | SE | | | | |
| 10 | | SE | NE | NW | | | | P2 |
| ⋮ | | | | | | | | |
| 13 | | | | | | | | P3 |

**B** Examples of swimming trajectories for NBN and Ctrl mice in each probe test.

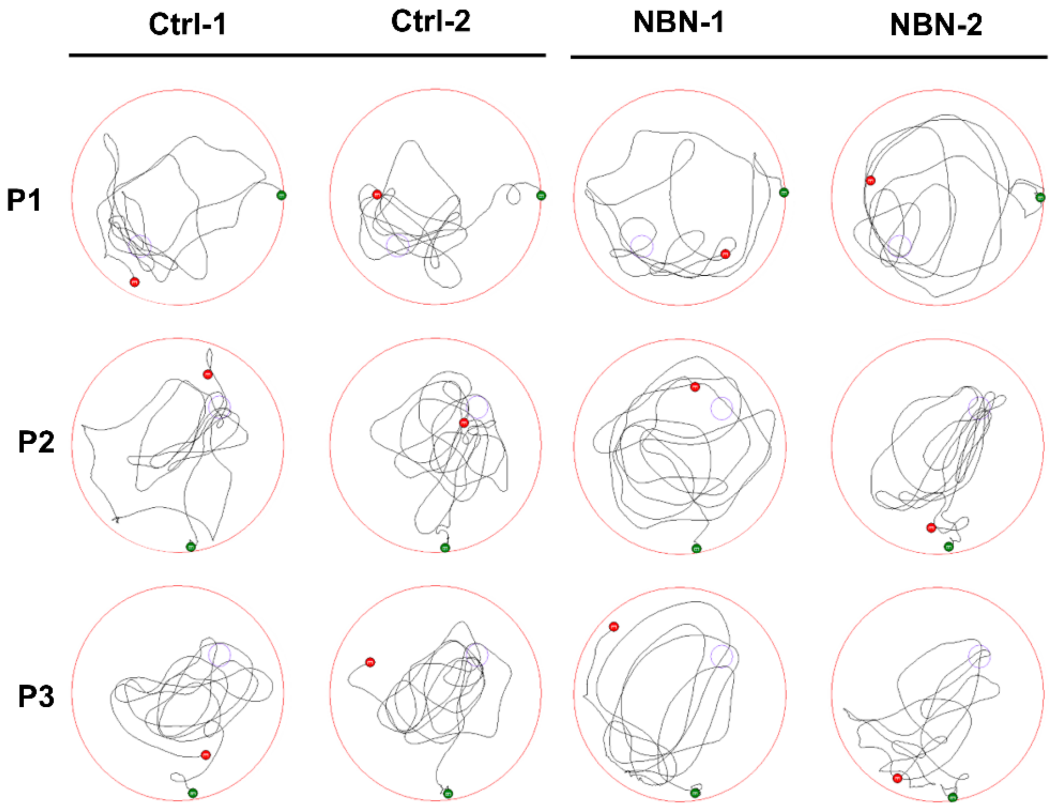


**Fig. S1.**

Starting Positoins and Swimming Trajectories in the Morris Water Maze.

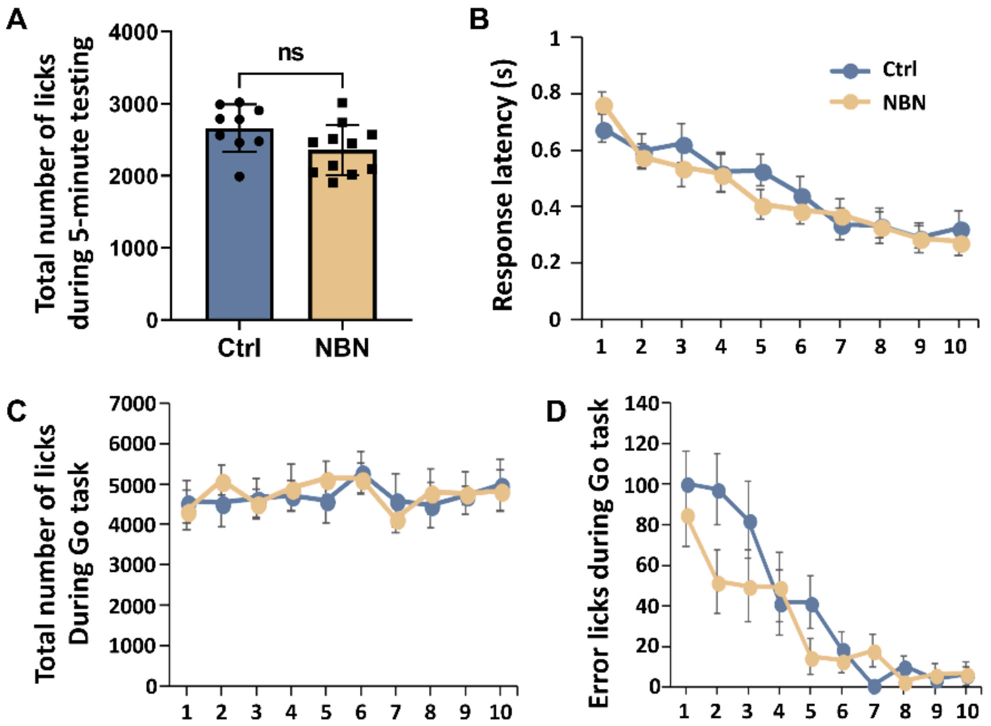


**Fig. S2.**

Additional Behavioral Assessment of Both Groups During Training and Go Task.

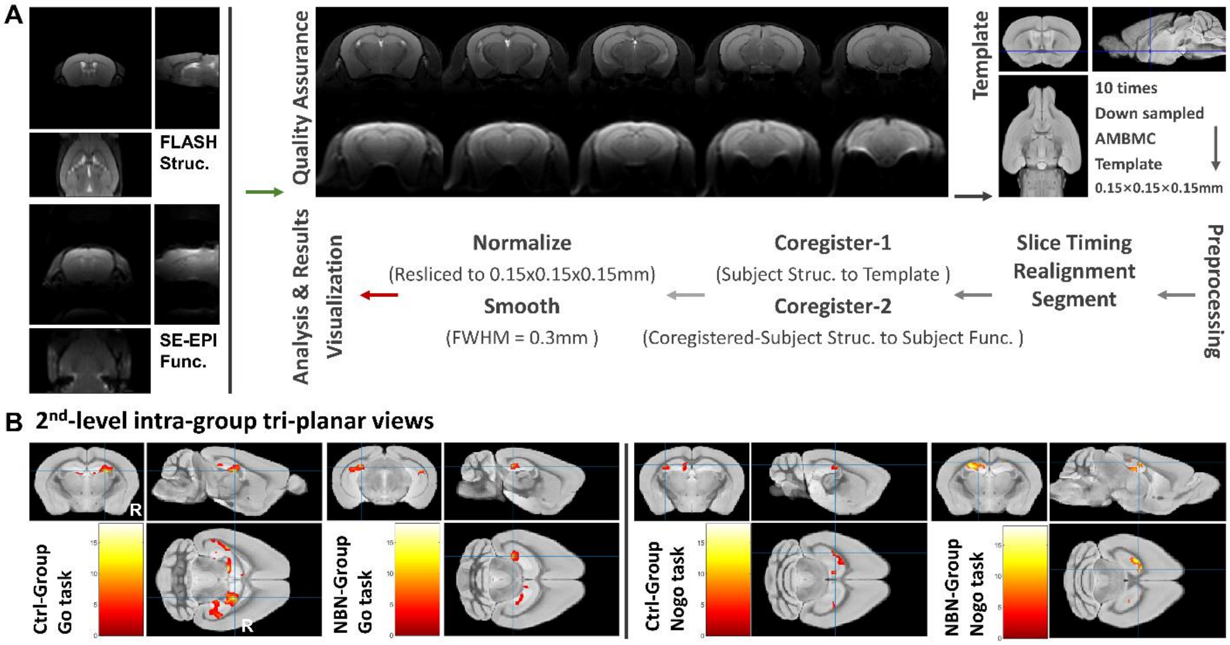


**Fig. S3.**

Protocol for Task-Based fMRI Analysis and Max T-Value Coordinates from Intra-Group Analysis of Go/Nogo Task.

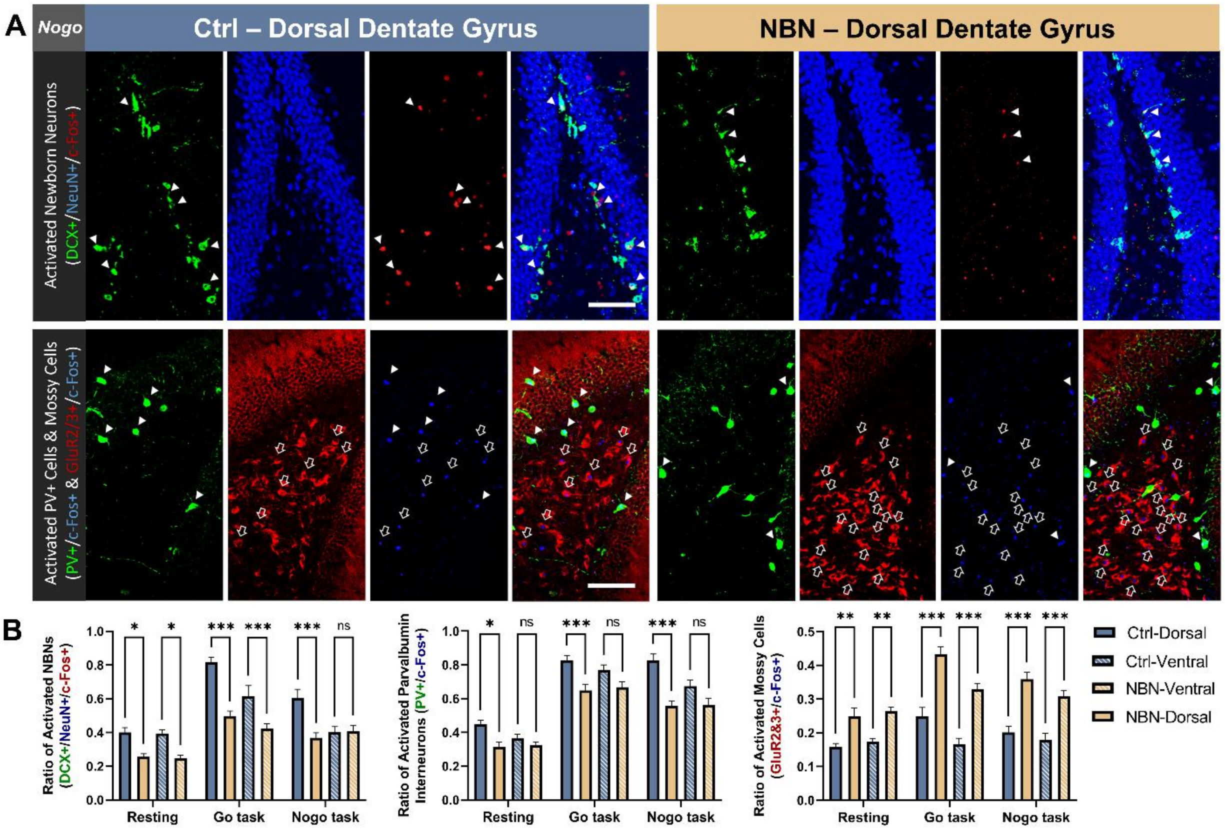


**Fig. S4.**

Post-Nogo Task Ventral Hippocampal Cell Activity Staining and Dynamic Changes Under Different Conditions.

**Table S1.**

Gene Sequence and PCR Conditions for NBN-TeTX Transgenic Mice.

| Gene Names | Primer sequences |
|---|---|
| Tg1: Nestin-CreERT2 (~340 bp) | 5'-TTCCGCTGGGTCACTGTCGCCGCTAC-3' (Fw) |
| | 5'-TAATCGCGAACATCTTCAGGTTCTGC-3' (Rv) |
| | **Reaction system composition** |
| | Template DNA (DNA solution): 1 µL |
| | 10 × PCR buffer: 10-fold dilution |
| | dNTP mixture: 0.2 mM |
| | Primer (Fw/Rv): 0.4 µM each |
| | Internal control primer: 0.4 µM each |
| | Taq: 0.3 units |
| | Deuterium-depleted water: up to 15 µL |
| | **Reaction condition** |
| | 95℃ 1min. |
| | 94℃ 30sec., 53℃ 45sec., 72℃ 1min.; × 32 cycles |
| | 72℃ 5min. |
| Tg2: αCamKII promoter-loxP-STOP-loxP-tTA (~500 bp) | **Primer sequences** |
| | 5'-CGCTGTGGGGCATTTTACTTTAG-3' (Fw) |
| | 5'-GGGTCCATGGTGATACAAGG-3' (Rv) |
| | **Reaction system composition** |
| | Template DNA (DNA solution): 1 µL |
| | 10 × PCR buffer: 10-fold dilution |
| | dNTP mixture: 0.2 mM |
| | Primer (Fw/Rv): 1.0 µM each |
| | Internal control primer: 0.5 µM each |
| | Taq: 0.45 units |
| | Deuterium-depleted water: up to 15 µL |
| | **Reaction condition** |
| | 94℃ 3min. |
| | 94℃ 30sec., 60℃ 30sec., 72℃ 30sec.; × 35 cycles |
| | 72℃ 2min. |
| Tg3: Tet Operator-TeTX (173 bp) | **Primer sequences** |
| | 5'-AAGTTCATCTGCACCACCG-3' (Fw) |
| | 5'-TCCTTGAAGAAGATGGTGCG-3' (Rv) |
| | **Reaction system composition** |
| | Template DNA (DNA solution): 1 µL |
| | 10 × PCR buffer: 10-fold dilution |
| | dNTP mixture: 0.2 mM |
| | Primer (Fw/Rv): 0.5 µM each |
| | Internal control primer: 0.5 µM each |
| | Taq: 0.45 units |
| | Deuterium-depleted water: up to 15 µL |
| | **Reaction condition** |
| | 95℃ 1min. |
| | 94℃ 20sec. |
| | Touch down over, (Initial temp. 65℃, Final temp. 60℃), 15sec. |
| | 68℃ 10sec.; × 10 cycles |
| | 94℃ 15sec., 60℃ 15sec., 72℃ 10sec.; × 28 cycles |
| | 72℃ 2 min. |
| **Internal positive control primer** | **Primer sequences** |
| for Tg1 (550 bp) | 5'-GTTAGCATTGAGCTGCAAGCGCCGTCT-3' (Fw) |
| | 5'-TGAGCGAGCTCATCAAGATAATCAGGT-3' (Rv) |
| for Tg2 & Tg3 (324 bp) | 5'-CTAGGCCACAGAATTGAAAGATCT-3' (Fw) |
| | 5'-GTAGGTGGAAATTCTAGCATCATCC-3' (Rv) |
